# B Cell Receptor Signaling Sets Boundaries for Selection of Broadly Neutralizing Antibody Functional Mutations

**DOI:** 10.64898/2026.08.13.744461

**Authors:** Ankita Singh, Kara Anasti, Elizabeth Van Itallie, Amanda Newman, Advaiti Pai Kane, Maggie Barr, Rob Parks, Sravani Venkatayogi, Ming Tian, Kevin Saunders, Rory Henderson, Derek W. Cain, Frederick W. Alt, Barton F. Haynes, Laurent Verkoczy, Kevin Wiehe, S. Munir Alam

**Affiliations:** Human Vaccine Institute, Duke University, Durham, NC, USA; Department of Medicine, Duke University, Durham, NC, USA; Department of Pathology, Duke University, Durham, NC, USA; Applied Biomedical Science Institute, San Diego, CA, USA; Howard Hughes Medical Institute, Program in Cellular and Molecular Medicine, Boston Children’s Hospital, Boston, MA, USA

**Keywords:** BCR, antigen, signaling, affinity, association rate, calcium mobilization, humanized knock-in, germinal center, B-cells, NGS, antibody, functional mutations, HIV-1, Envelope, neutralization

## Abstract

B cell signaling is required for germinal center (GC) selection and synergizes with T cell-help leading to differentiation into effector cells and a protective antibody response. Here, we studied the relationship between B cell signaling and the selection of functional mutations in a humanized mouse model of a CD4 binding-site specific HIV-1 broadly neutralizing antibody precursor. While BCR signaling increased with antigen affinity, immunization-induced frequency of a functional mutation was inversely related to affinity and favored a gain in association rate. Antigen-specific GC B cells and key mutation frequency were higher in the mid-affinity (0.5 – 5μM) than in either the higher or lower affinity group, and were consistent with the significantly higher serum neutralization titers in the mid-range group. Our studies show that BCR signaling imposes boundaries (upper/lower) for selection of antibody functional mutations and support a “Goldilocks Zone” model that defines the favored BCR signaling strength for GC selection.

## Main

The germinal center (GC) is a privileged microanatomical environment for naïve B cells that have encountered an antigen to undergo somatic hypermutation (SHM) and cell fate decisions including survival, proliferation and differentiation into effector cells (plasma/memory)^1–3^. Within the GC, cycling of B cells between light (LZ) and dark zone (DZ)^4^ and SHM-dependent selection of mutations drive the affinity maturation process that results in the development of high affinity antibodies and effective humoral immunity^5–7^. According to the affinity model of GC selection, B cell receptor (BCR) antigen affinity is a key determinant for B cell survival and for GC B cell differentiation into memory or plasma B cells^8^. Recent studies have shown that B cell signaling is required for both survival and priming of GC selection^9,10^. Importantly GC B cell signaling synergizes with cognate T cell help for positive selection and differentiation^1,6^. However, the range of BCR signaling strength and whether there are upper/lower thresholds that support GC selection remain unclear. Although weak affinity GC B cells are more likely to undergo apoptosis^11^, the lower affinity threshold is not clearly defined. In both early and late GCs, B cells having antigen affinity of a wide range (μM to nM) and even those with no measurable antigen affinity have been reported^12^. In the absence of competition with high affinity B cells or when the T cell help is abundant, low affinity B cells can enter GCs and undergo affinity maturation^13–15^. At low precursor frequency or when subject to competition, relatively higher affinity outperforms weaker affinity B cells^12,14,16^. At the other end of the affinity spectrum, high affinity binding can lead GC B cells towards extrafollicular/plasmacytic rather than GC differentiation^17–19^. Therefore, too high an affinity might not lead B cells towards a sufficiently prolonged SHM process, rather prematurely ending the GC program and consequently resulting in a dearth of SHM-derived functional mutations. It follows from the above, that there is likely a lower and upper limit of antigen affinity-dependent BCR signaling, yet not clearly defined, that governs the length of LZ-DZ cycling and selection of affinity-enhancing mutations. Recently, we reported that the strength of BCR signaling is dependent on antigen binding association rate and a threshold antigen-BCR half-life^20^, and therefore, BCR signaling is coupled to antigen internalization and consequently to T cell help. However, the relationship between BCR signaling strength dependence on antigen binding association rate and selection for affinity-enhancing and/or functional mutations is not clearly understood.

For a protective humoral response to viral pathogens, the selection of functional mutations is critical for the development of antibodies with higher affinity as well as plasticity in epitope recognition, as in the case of broadly neutralizing antibodies (bnAb) that target conserved sites on the HIV-1 Envelope (Env) protein^21,22^. The numerous HIV bnAbs that have been isolated display common features that include high mutation frequency, often with insertion-deletions (in-dels)^23^, long CDRH3s, and key functional mutations that have a low probability of occurrence (improbable mutations) due to biases in the activation-induced cytidine deaminase (AID) catalyzed activity during the SHM process^24^. The conserved CD4 binding-site (CD4-bs) on HIV Env is a key target for a class of bnAbs that mimic CD4 binding for their ability to neutralize. Among the most potent CD4-bs bnAbs that utilize the variable heavy chain 1-46 (VH1-46), the CH235 lineage presents a key bnAb model due to high viral neutralization breadth (90%), lack of indels, and normal CDR3 lengths^25,26^. Critical improbable and functional mutations in the CH235 bnAb lineage have been extensively characterized and several Env trimer gp140 proteins that bind to the inferred CH235 unmutated common ancestor (CH235.UCA) antibody have been designed^27–29^. Env trimers that bind CH235.UCA with a relatively high-affinity can induce selection of key functional mutations in a humanized knock-in mouse model (CH235.UCA KI) that expresses B cells with BCRs with CH235.UCA VH/VL specificity^28,30^. In this study, to define the range of affinity/kinetic rates that is permissible for selection of key functional mutations in the CH235 bnAb lineage, we performed immunization studies in the CH235.UCA KI mice using Env gp140 trimers that bind with a wide range of affinities/kinetic rates and trigger varying strength of BCR signaling. Here, we report (i) the relationship between BCR signaling strength and the selection of critical mutations; (ii) define the antigen binding parameter that drives high frequency selection of a key mutation, and (iii) define the lower and upper limits of BCR signaling that govern selection of antibody functional mutations and viral neutralization potency.

## Results

### HIV-1 gp140 trimer binding affinities and kinetic rates to CH235.UCA antibody

In this study, we selected a panel of five CH505 gp140 Env trimer proteins that bound to the CD4 binding-site (CD4-bs) specific CH235 bnAb unmutated common ancestor antibody, CH235.UCA (**Fig. 1A; Table S1**). The trimers are numbered (#1-5) by decreasing affinity to CH235.UCA with 1 having the highest and 5 the weakest affinity. CH235.UCA Fab bound to the trimers with affinities (K_D_) from μM (10^-6^ M) to nM (10^-9^ M), and with distinct association (k_a_) and dissociation (k_d_) rates (**Fig. 1A**). The two highest affinity trimers (#1-2) both bound with fast k_a_ (101 and 53.7 x10^3^ M^-1^s-1, respectively) and with similar and slow k_d_ (10.7 and 9 x10^-4^ s-1, respectively). The three moderate and weak affinity trimers (#3-5) had similar k_d_ (101 - 336 x10^-4^ s-1) but bound with k_a_ values that differed by 20-fold (0.9 - 18.2 x10^3^ M^-1^s-1) (**Fig. 1A, S1**). Thus, the increase in trimer affinity (#5 to 1) was predominantly due to a progressive increase in k_a_ (association rate/on-rate). The selected panel of trimeric proteins provided a wide range of affinities, spanning three orders of magnitude (K_D_= 23.5 μM to 10.6 nM) and with differing kinetic on-rates, to study *in vitro* B cell activation and *in vivo* immunization studies in a knock-in mice (KI) model expressing BCRs with specificity of CH235.UCA.

**Figure 1.**
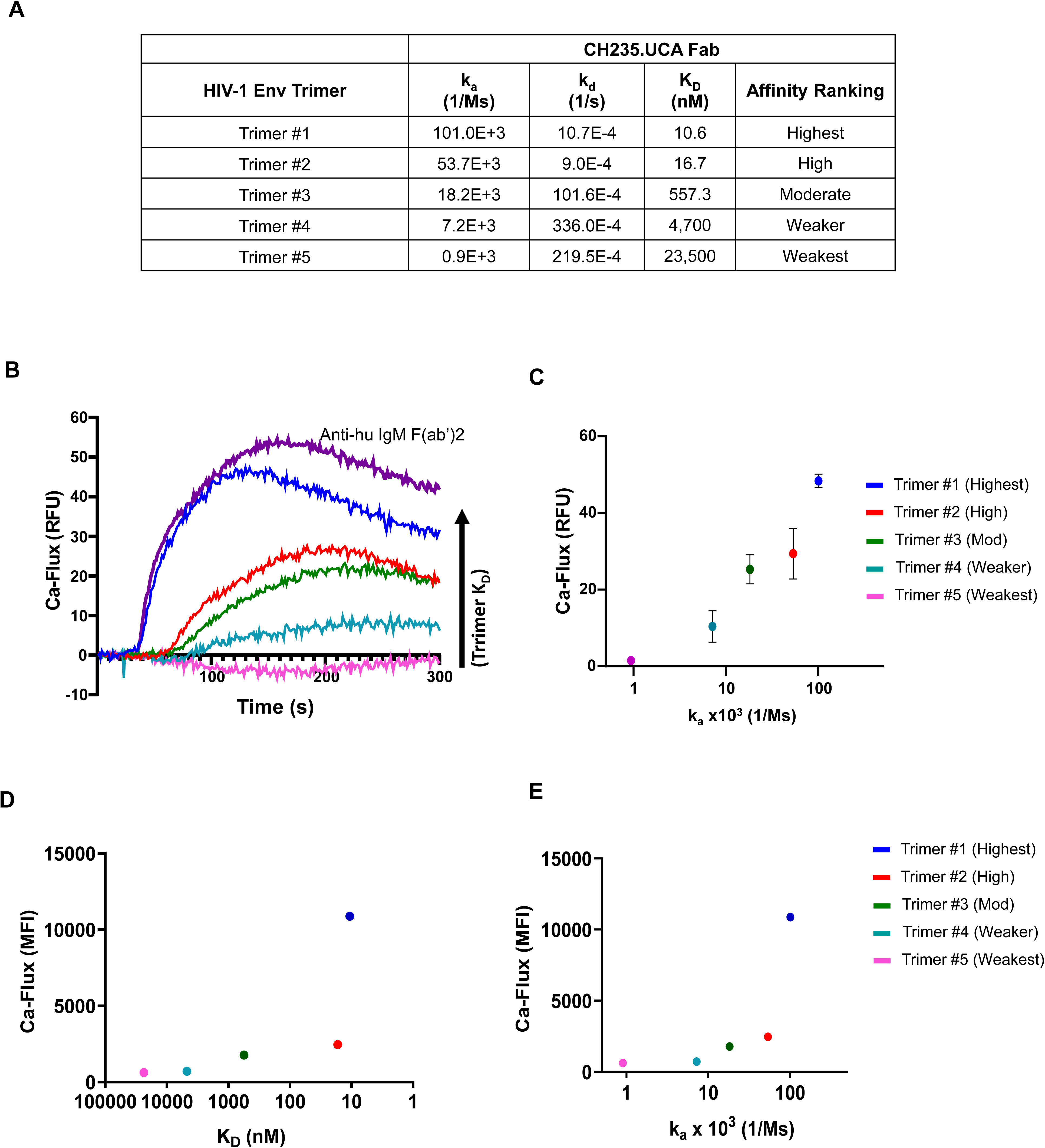
BCR signaling strength and trimer affinity. (**A**) Surface plasmon resonance (SPR) measured kinetic rates (k_a_, k_d_) and affinities (K_D_) of CH235.UCA Fab to 5 different HIV-1 Env CH505 trimer proteins ranked by affinity. The affinities of trimers# 1, 4, and 5 with CH235.UCA Fab were previously reported^29,30^. **(B)** Calcium flux (Ca-flux) responses in CH235.UCA-IgM Ramos cells induced with trimer proteins. CH235.UCA-IgM Ramos cells were treated with trimers #1 (dark blue), #2 (red), and #3 (green) at 100nM while #4 (light blue) and #5 (pink) trimers were each tested at 250nM. Calcium responses were monitored for 5 minutes after treatment with trimers. Activation with the positive control anti-human IgM F(ab)_2_ control cross-linking antibody is shown (purple). **(C)** Maximum Ca-flux responses induced by trimers in CH235.UCA-IgM Ramos cells (y-axis) versus trimer association rates (k_a_, 1/Ms) (x-axis) to CH235UCA Fab. Ca-flux responses are calculated as relative fluorescence unit (RFU). **(D)** Ex-vivo Ca-flux response of CH235.UCA knock-in (KI) splenic B cells versus trimer protein affinity (K_D_) to CH235.UCA Fab. **(E)** Ex-vivo Ca-flux response of CH235.UCA KI B cells plotted versus trimer association rate (k_a_). The ex-vivo data are shown as median fluorescence intensity (MFI).The x-axis shows K_D_ and k_a_ on a logarithmic scale. F(ab’)2-anti-mouse IgM (µ chain) was used as the positive control for the ex-vivo study that showed mean Ca-flux peak response at 10429.33 MFI. The M5 G458Y N280D trimer was used as the negative control and the peak response was observed at 512 MFI. The N280D abrogates binding of CD4-bs and neutralization and it had no measurable affinity. The baseline peak was used for background subtraction. The details of trimers used in the present study are given in Table S1.

### CH235.UCA IgM BCR signaling strength is dependent on antigen binding association rate

To measure *in vitro* B cell activation to the panel of five trimers that bind to CH235.UCA Fab with different kinetic rates and affinities (**Fig. 1A**), we developed a Ramos B cell line expressing IgM BCRs with V_H_/V_L_ sequences of CH235.UCA (CH235.UCA IgM cells) (**Fig. S2**). The CH235.UCA IgM cells express functional IgM BCRs and co-receptors and gave strong calcium flux response (Ca-flux), an early indicator of B cell activation, following stimulation with the BCR crosslinking anti-human IgM F(ab’)_2_ control antibody (**Fig. 1B**). We observed differential Ca-flux responses on CH235.UCA IgM cells when stimulated with the selected trimers (#1-5) and the peak Ca-flux response increased with increasing affinities of the trimers (**Fig. 1B**). When comparing the two highest affinity trimers (trimer #1-2) that had almost identical k_d_ (off-rate) rates, the trimer with the fastest k_a_ (#1) gave markedly higher Ca-flux response that was the strongest among all trimers (**Fig. 1B**). Notably, for the remaining three trimers (trimer #3-5) that bound with a wide range of affinity but similar k_d_, the Ca-flux response magnitude increased with increasing k_a_ (on-rate) values (**Fig. 1B-C**). The phosphorylation profile of the kinases (syk, btk, blnk) that comprise the Ca-flux signalosome complex was similar to the Ca-flux responses to the trimers (**Fig. S3**). Thus, the highest affinity trimer induced high levels of phosphorylation of the three phosphokinases, while the weaker affinity trimers with μM K_D_ induced low levels of phosphorylation explaining their inability to induce strong Ca mobilization (**Fig. 1B, S3**). Overall, the magnitude of Ca-flux responses showed a positive relationship with the on-rate (k_a_) of the trimers to CH235.UCA Fab (**Fig. 1C, E, S4A-B**).

Next, we measured *ex-vivo* Ca-flux response in splenic B cells isolated from the CH235.UCA V_H_/V_L_ knock-in (KI) mice (**Fig. S5A-D**). As observed with the CH235.UCA IgM Ramos cells, the CH235.UCA KI B cells showed a similar affinity/k_a_ relationship with Ca-flux responses with the panel of trimers (**Fig. 1D-E, S4C, S5B-C**). Thus, in both CH235.UCA IgM Ramos B cells and CH235.UCA KI primary B cells, we observed that the strength of early B cell signaling increases with binding association rates and these results are consistent with our previous studies on a different class of CD4-bs specific BCR-expressing B cells^20^.

### Magnitude of immunogen-specific sera IgG titers is BCR signaling dependent

To investigate *in vivo* the role of antigen binding affinity/rates and BCR signaling potential in naïve B cell activation and selection of functional mutations, we performed two immunization studies in the CH235.UCA KI mice that harbor B cells expressing BCRs with the pre-rearranged V(D)J rearrangements of the CH235.UCA antibody^27^. In the first study, we had three groups of mice each immunized with either the high (K_D_=16.7nM, trimer #2), moderate (K_D_= 557.3nM, trimer #3), or weaker (K_D_= 4,700nM, trimer #4) affinities trimers. In the second study, we had two groups immunized with either the highest (K_D_= 10.6nM, trimer #1) or the weakest (K_D_= 23,500nM, trimer #5) affinity immunogens. The immunogen dose (25ug), adjuvant (5ug SMNP), and sequential immunization regimen were the same in the two studies, and the aggregate results are thus presented in **Fig 2**.

**Figure 2.**
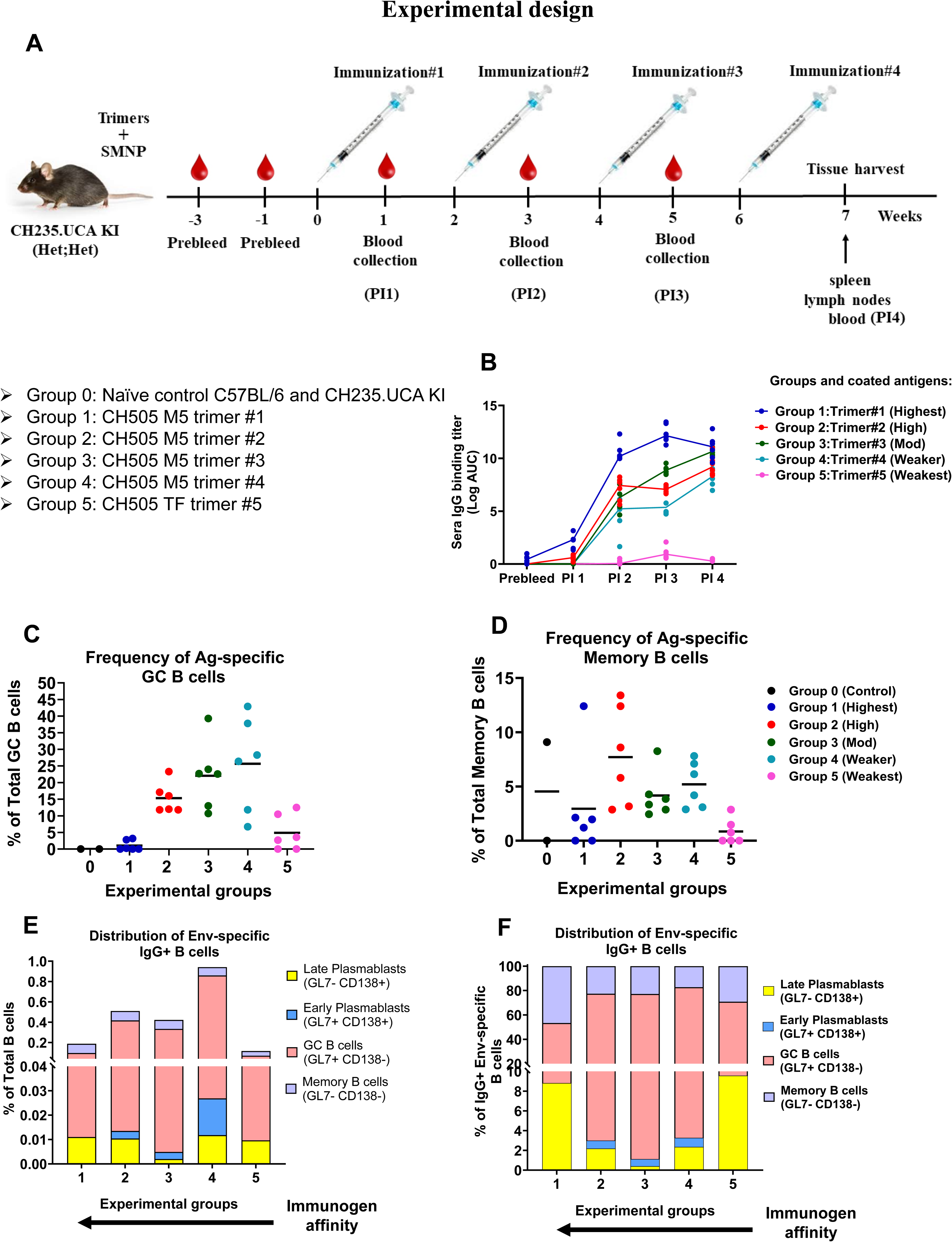
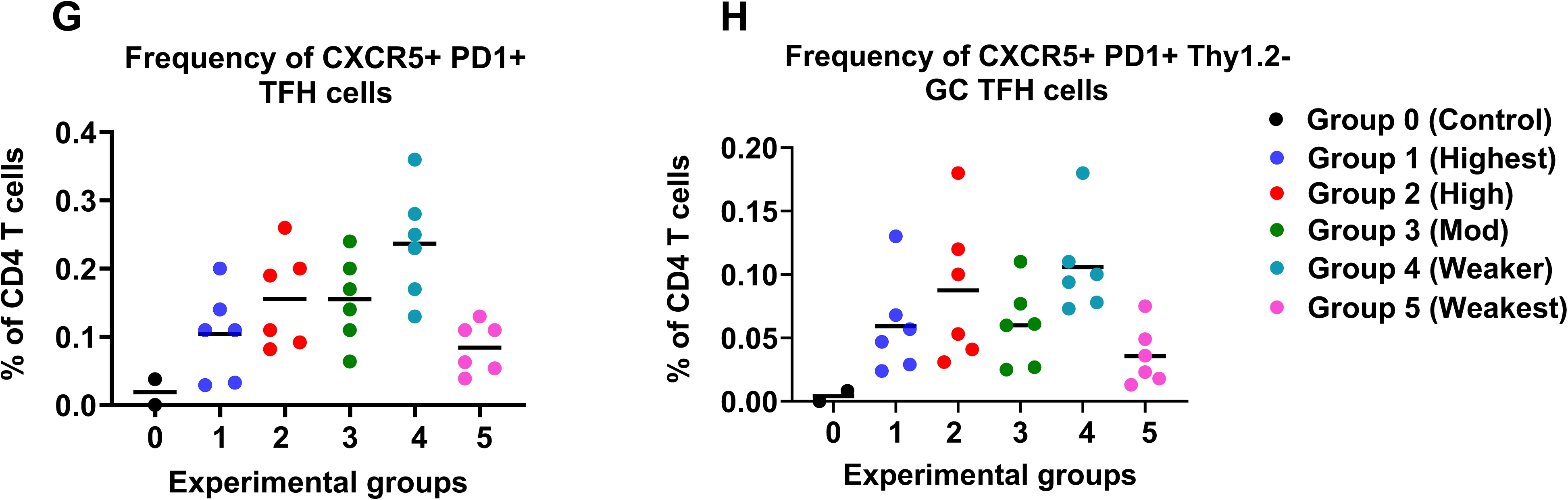
Sera antibody titers and cell phenotypes in immunized CH235.UCA KI mice. **(A)** Experimental design of immunization in CH235.UCA KI (heterozygous) mice. Sequential immunization was performed four times at two-week intervals. Mice were injected on leg intramuscularly with 25µg trimer and 5µg SMNP adjuvant. Blood sample draws and tissue harvests were done at indicated time points. (**B)** Kinetics of sera antibody binding titer (logAUC) in each experimental group. Antigen-specific sera IgG binding titer (y-axis) versus different post immunization (PI) time points (x-axis). **C-D** Lymph node antigen-specific B cell phenotyping in various experimental groups **(C)** Frequency of antigen-specific GC B cells among total GC B cells (y-axis) versus experimental groups 0-5 (x-axis) **(D)** Frequency of antigen-specific memory B cells among total memory B cells (y-axis) versus experimental groups 0-5 (x-axis). **E-F** Distribution of lymph node class-switched antigen-specific B cells into plasmablast, memory, and GC compartments. **(E)** The plot shows the frequency of IgG+ Env+ B cells in various compartments as a percent of total B cells (y-axis) plotted versus experimental groups (x-axis). **(F)** The plot shows the frequency of IgG+ antigen-specific B cells in different compartments as a percent of class-switched IgG+ Env-specific B cells (y-axis) versus immunized experimental mice (x-axis). **G-H** Lymph node T cell phenotypic analysis in each experimental group **(G)** Frequency of T follicular helper (TFH) cells among CD4 T cells (y-axis) versus experimental groups 0-5 (x-axis). **(H)** Frequency of GC TFH cells among CD4 T cells (y-axis) plotted versus experimental groups 0-5 (x-axis). Each dot represents an individual mouse and bar indicates the mean value. Experimental groups #1-5 were immunized with trimers #1-5, respectively. Naïve wild-type C57BL/6 and CH235.UCA knock-in mice were kept as control group 0. Statistical analysis was done using exact Wilcoxon test. *P* < 0.05 was considered statistically significant. Relative proportion of GC B cells to plasmablasts is given in Table S2. *P*-values for ELISA and B and T cell phenotyping are given in Tables S3 and S9-S11 respectively.

To determine the antigen-specific IgG antibody titers, sera samples were collected at pre- and post-immunization (PI) time points (**Fig. 2A, S6**) from immunized mice in each experimental groups and were examined for binding to each of the trimer immunogens by ELISA. In each group, low binding titers were detected after the first immunization (PI 1) and peak titers were observed after the third (PI 3) or fourth (PI 4) immunization (**Fig. 2B, S6A-E, Tables S3-S5).** Sera from each immunogen groups showed reactivity to each of trimer proteins in the studied panel (**S6A-E, Tables S4-S8**). Immunogen-specific IgG titers in PI4 samples showed higher titer in the highest affinity group (group1, 10.6nM) compared to groups 2, 4, and 5 (group 1 > 2 p=0.026, group 1 > 4 and 5 p=0.0022) whereas lowest titer were in the weakest affinity group (group 5, 23.5µM) compared to groups 1-4 (10.6nM – 4.7µM) (groups 1-4 > 5 p=0.0022) (**Fig. S6F, Table S3**). However, binding to the weakest affinity trimer #5 remained minimal across all immunogen groups (**Fig. S6E, S6E-G, Tables S4-S5, S8**), likely reflecting the relatively weak affinity of the induced antibodies to trimer #5 and not due to low sera IgG antibody induction as evident in the observed strong titers when assayed against the higher affinity immunogen trimers (**Fig. S6A-D**). Notably trimer #5 binds well to the affinity matured CH235 lineage intermediate antibodies (I59-I39, 12.7- 5.3nM), but with much weaker affinity to the first intermediate I60 (25.6μM)^28,30^ (**Fig. S7**), indicating a likelihood of limited affinity maturation in group 5 animals. Overall, the serology analysis showed that each immunogen induced sera IgG responses, and the magnitude of immunogen-specific IgG titers (**Fig. 2B**) showed a trend that reflected the strength of BCR signaling (**Fig. 1B, S6F**) triggered by the five trimer immunogens.

### Higher frequencies of antigen-specific IgG+ GC B cells in response to immunogens within an affinity range

As in our previous study^30^ and due to the enhanced antigen lymphatic flow and robust B cell responses with the administered adjuvant^31^, we collected lymph nodes (LNs) to study the B cell responses in the immunized CH235.UCA mice. We compared LN trimer-specific B cell phenotypes and frequencies of either germinal center or memory B cells in each of the immunized groups (**Fig. S8A-C, S9A-D**). Among total GC B cells, mice immunized with either the weakest (group 5, mean 4.88%) or the highest affinity (group 1, mean 1.04%) trimers showed the lowest mean frequency of antigen-specific GC B cells (**Fig. 2C, Table S9**). The mean frequency of antigen-specific GC B cells was higher (mean 15.33% - 25.65%) in the immunized groups within a wide affinity range (groups 2-4, 4.7μM to 16.7nM) (groups 2, 3, and 4 > 1 p=0.0022; groups 2, 3, and 4 > 5 p=0.0130, 0.0043, 0.0152 respectively). However, the mean GC B cell frequency was highest in the weaker affinity group 4 mice (mean 25.65%), indicating a bias in favor of the lower immunogen affinity (4.7μM) but not the weakest affinity (23.5μM) (**Fig. 2C, Table S9**). While the antigen-specific memory B cells were relatively lower in all five groups, we observed a trend in which the mean frequency was lowest in the two groups (1 & 5) outside the favored affinity boundaries in the GC B cell compartment (**Fig. 2D, Table S10**). Among total B cells, the frequency of IgG+ antigen-specific B cells was also lower in groups 1 and 5 compared to other immunized groups (**Fig. S9E**).

Next, to examine the effect of immunogen affinity on B cell differentiation into GC, memory, and plasmablast (PB) B cells, we studied the distribution of lymph node antigen-specific IgG B cells into each of the above compartments (**Fig. S8E**). Among total B cells, the antigen-specific IgG+ GC B cell phenotype was predominant in all groups (**Fig. 2E**). However, the relative proportion of GC B cells to late PB was higher in groups 2-4 (group 2: 38.8, group 3: 163.3, group 4: 70.6) when compared to groups 1 (7.7) or 5 (6.2) (**Fig. 2E, Table S2**). Of the IgG+ antigen-specific B cells, similar results were observed where the proportion of late PBs was substantially higher in the lowest and highest affinity groups than in the moderate affinity groups (**Fig. 2F, Table S2**). These results suggest that both groups 1 (highest affinity) and 5 (weakest affinity) have a relatively higher plasmacytic response than those in the moderate affinity groups (2-4).

In the analysis of the frequency of T-follicular helper (TFH) cells as a measure of B cell helper function (**Fig. S8D**), we observed a similar trend as in the GC B cell frequency. The mean frequency of CXCR5+PD1+ TFH cells progressively increased with decreasing affinity except for the weakest affinity group 5 (group 4 > 1 p=0.0130; groups 3 and 4 > 5 p=0.0390, 0.0043 respectively) (**Fig. 2G, S9, Table S11**). The frequency difference of PD1+ GC TFH cells was relatively less apparent in the immunized groups 1-4, but the mean % GC TFH cells was the lowest in group 5 (group 4 > 1 p=0.0411; group 4 > 3 p=0.0433; group 4 > 5 p=0.0043) (**Fig. 2H, S9, Table S11**). The frequency of TFR cells remained low in all groups but with groups 1 and 5 showing relatively higher mean % values (**Fig. S9H, Table S12**). These results, therefore, show that both higher frequency of lymph node GC B (antigen-specific) and TFH (CXCR5+PD1+) cells are observed in response to immunogens that are within an antigen affinity range (∼20nM to 5μM) but are significantly lower outside this boundary at both ends of the affinity spectrum.

### BCR signaling strength sets boundaries for induction of autologous neutralizing antibody responses

To evaluate the neutralizing antibody titers elicited in the immunized mice, we collected PI4 sera samples from each group of the immunized mice and measured sera ID_50_ titers in the HIV pseudovirus neutralization assay. We tested neutralization against two autologous pseudoviruses, CH505TF.M5 and CH505TF. M5.G458Y. The latter with the affinity enhancing G458Y mutation is neutralized by CH235.UCA, while the former without G458Y is the more difficult to neutralize autologous viral strain^28^. Importantly, sera from each of the immunized groups showed strong IgG binding responses to soluble immunogen trimers (**Fig. 3A-B**) that matched the Env protein mutations (M5 or M5+G458Y) expressed on the pseudoviruses used in the neutralization assay.

**Figure 3.**
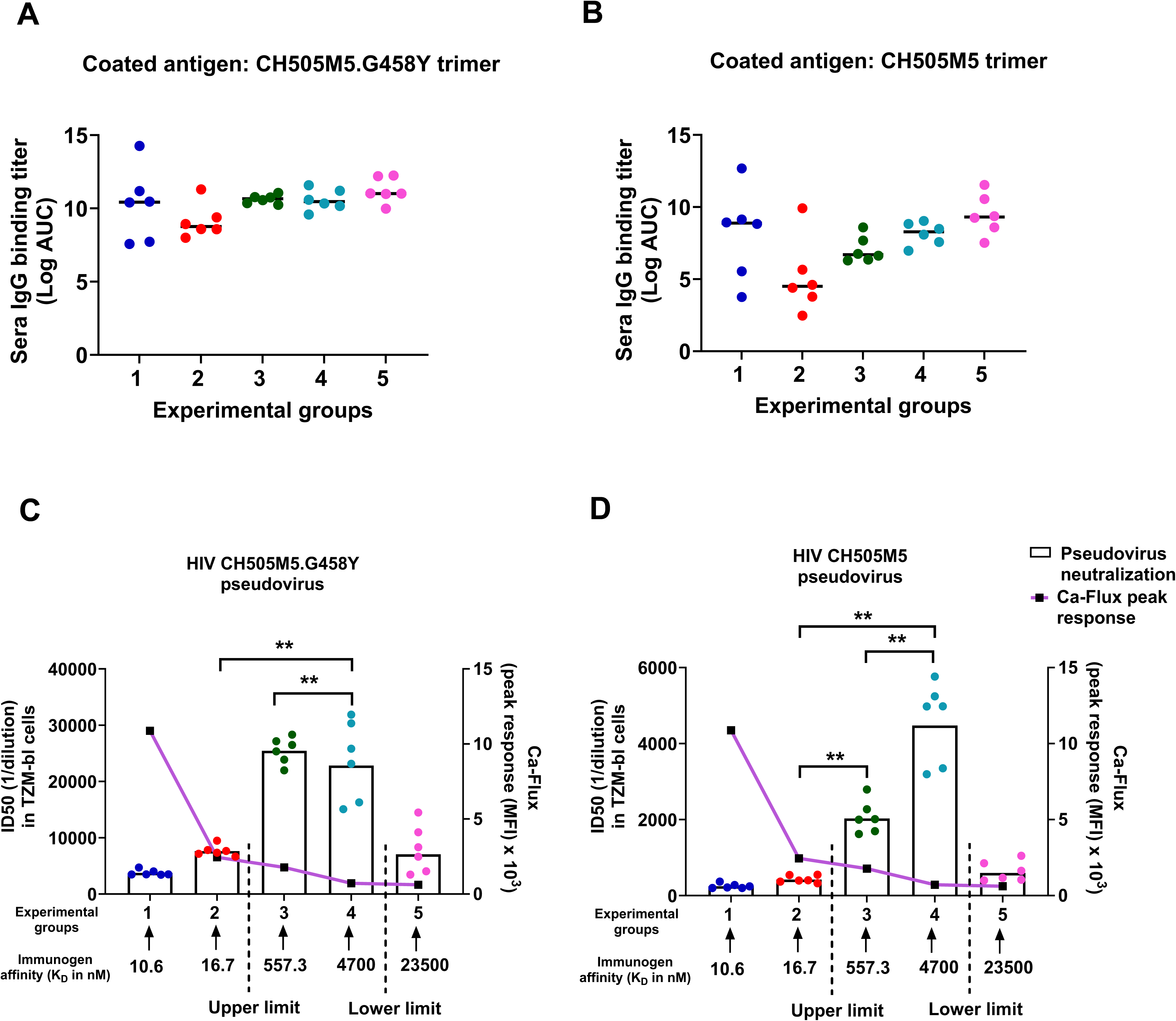
Sera binding titers and pseudovirus neutralizing titers in immunized mice. **(A-B)** ELISA measured antibody binding titers in each group at PI4. Sera IgG binding titers to **(A)** the moderate affinity trimer (CH505M5.G458Y) and **(B)** the weaker affinity trimer (CH505M5). Binding titers are shown as log area under the curve (AUC). **(C-D)** ID50 neutralization titer at PI4 in a pseudovirus neutralization assay. **(C)** Sera ID50 neutralization titers against autologous CH505TF.M5 G458Y pseudovirus (y-axis1) and ex-vivo Ca-flux peak responses (y-axis2) plotted versus experimental groups 1-5 (x-axis). **(D)** Sera ID50 neutralization titers against autologous tier 2 CH505TF.M5 pseudovirus (y-axis1) and ex-vivo Ca-flux peak responses (y-axis2) plotted versus experimental groups 1-5 (x-axis). Titers are represented as reciprocal sera dilution required to neutralize 50% of viral replication (ID50). The solid line connects the Ca-flux peak values for each experimental group. K_D_ (nM) values of the trimeric immunogens for each immunized group are given on the x-axis. Dotted lines indicate upper and lower affinity limits for potent neutralization. Each symbol represents an individual mouse, and bars indicate the geometric mean. Statistical analysis between groups was performed using exact Wilcoxon test and statistically significant differences are denoted by asterisk (*P < 0.05, **P < 0.01). MFI=median fluorescence intensity. P-values for sera binding titers and neutralization assay are given in Tables S3 and S13 respectively.

The sera neutralization from each group was sensitive to the CD4 binding site (CD4-bs) specific knockout N280D mutation^28,29^ (**Fig. S10**), indicating induction of antibodies against the targeted CD4-bs epitope. All five groups of immunized mice sera showed neutralization, although of varying potency (ID_50_), against the CH505.M5 G458Y pseudovirus (**Fig. 3C**). The mean (geometric) ID_50_ neutralization titers were significantly higher in the moderate and weaker affinity groups 3 and 4 (ID_50_=25,439 and 22,834 respectively) when compared to either the two higher affinity groups (1 & 2, ID_50_=3,749 and 7,635 respectively) (groups 3 and 4 > 1 and 2 p=0.002) or the weakest affinity group 5 (ID_50_=7,051) (groups 3 and 4 > 5 p=0.002) (**Table S13**). When tested against the pseudovirus without the affinity enhancing G458Y mutation (CH505TF.M5), the same neutralization trend was observed; higher potency in groups 3 and 4, while both the higher (1 & 2, mean ID_50_=246 and 423 respectively) and the weakest (group 5, mean ID_50_=597) affinity group sera gave markedly lower ID_50_ titers (groups 3 and 4 > 1, 2, and 5 p=0.002) (**Fig. 3D, Table S13**). Neutralization ID_50_ of the tier 2 CH505TF.M5 pseudoviruses was significantly higher in group 4 (mean ID_50_=4,474) when compared to group 3 (mean ID_50_= 2,030) (group 4 > 3 p=0.002). No neutralization was observed against the tier 2 CH505TF pseudovirus that has neither the M5 nor the G458Y mutations (**Fig. S10**).

Overall, the sera neutralization results show that the titers are significantly higher in immunogen affinity groups within a boundary (lower and upper, dashed lines in **Fig. 3C-D**) and are not dependent on the magnitude of sera IgG binding titers to trimers that matched the Env expressed on the two peudoviruses **(Fig. 3A-B)**. Importantly, the boundaries can be defined by trimer affinity-dependent BCR signaling strength (**Fig. 1C-E**). The lower boundary of immunogen affinity (∼20μM) corresponds to an on-rate threshold (apparent k_a_ <10^3^ M^-1^s^-1^, increasing 8-fold between trimer #5 and 4), while the upper boundary is observed at an apparent affinity ceiling (∼20nM) with an increase in on-rate (3-fold, trimer #3 vs 4) and an order of magnitude slowing of the off-rate (k_d_ ∼10^-4^ s^-1^, 11-fold trimer #3 vs 2) (**Fig. 1A, 3**). At the far end of the affinity spectrum, the highest affinity trimer #1 has the fastest on-rate (10^5^ M^-1^s^-1^), resulting in the strongest BCR signaling (**Fig. 1C-E**), but that severely limited the ability of the immunogen to induce high titers of neutralizing antibody responses (**Fig. 3C-D**). Thus, there is a narrow affinity range (∼5μM to 0.5μM) and a corresponding optimal BCR signaling strength that is most favorable for induction of antibody responses with high neutralizing titers. We, therefore, hypothesized that too strong or too weak BCR signaling would result in antibody responses with lower frequency of GC selection of antibody functional mutations that are required for viral neutralization.

### Selection of functional mutations favors weaker BCR signaling

The observed differences in the GC B cell frequency and the bias in neutralization titers led us to hypothesize that there are qualitative differences in the antibodies induced in the five immunized groups. Therefore, we reasoned that the efficiency of selection of antibody functional mutations might be different in response to immunogens with differing BCR signaling potential. To address the above hypothesis, we examined whether immunization in the CH235.UCA KI mice will induce selection of key functional antibody mutations that were identified during the affinity maturation of the CH235 lineage^24,26,28^. Thus, we performed next-generation sequencing (NGS) of heavy and light chain rearrangements (V_H_DJ_H_ and V_L_JL) on unsorted splenic B cells from each of the immunized mice in the immunogen groups 2-5 (affinity range 16.7nM to 23.5μM). The highest affinity (10.6nM) group 1 mice were not included in the NGS analysis because neutralization titers in group 2 (16.7nM) included in the NGS analysis were similarly low (**Fig. 3C-D**).

To identify the key functional mutations that occurred in the maturation of the CH235 lineage, we compared the VH amino acid sequences in the inferred UCA and proximal intermediate antibodies (I60, I59) in the CH235 bnAb lineage maturation pathway^30^ (**Fig. 4A**). The first intermediate I60 has eight amino acid differences of which five are improbable (**Fig. 4A**). There are an additional two amino acid differences between the first (I60) and second (I59) intermediates, one of which is improbable (**Fig. 4A**).

**Figure 4.**
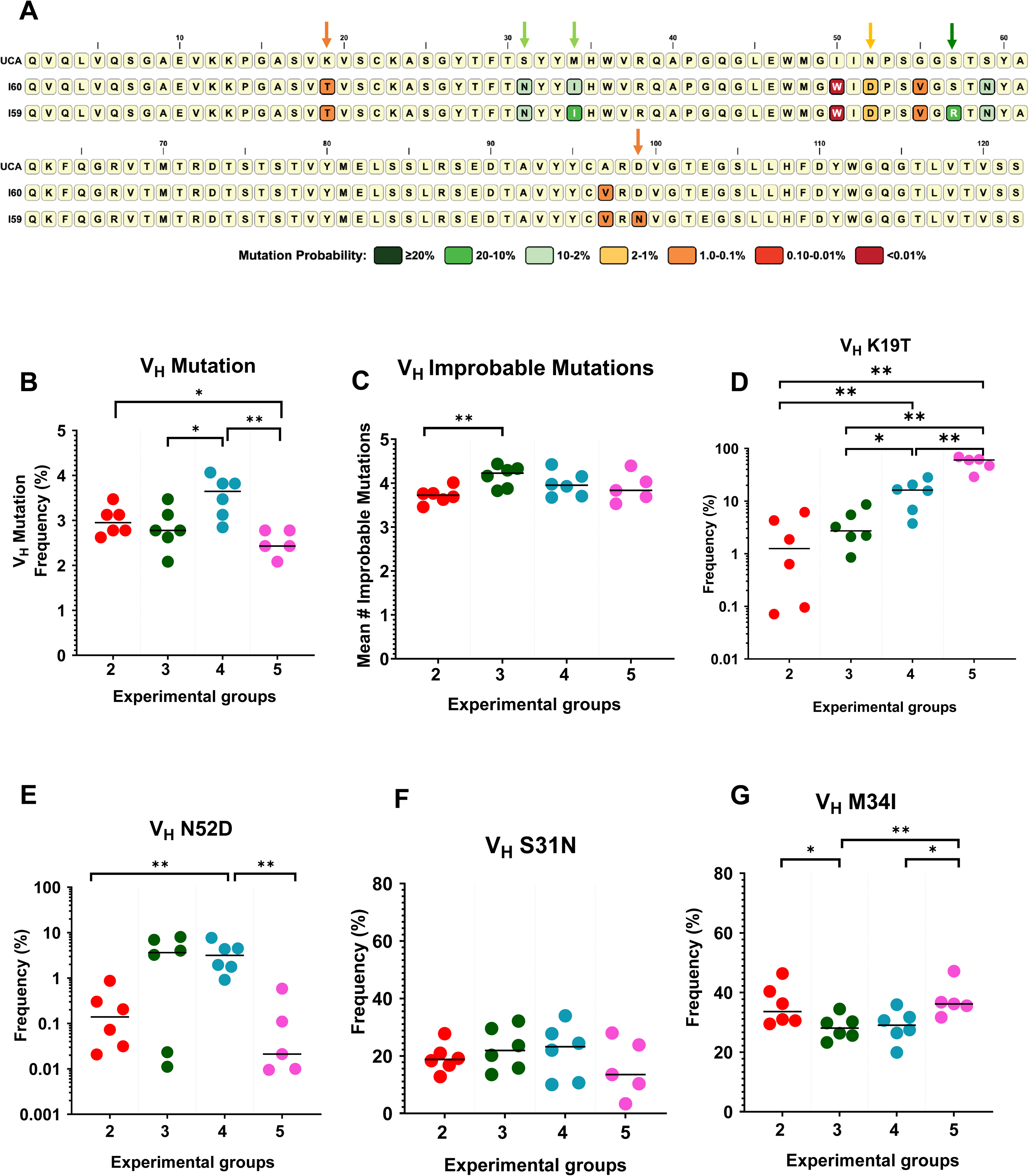
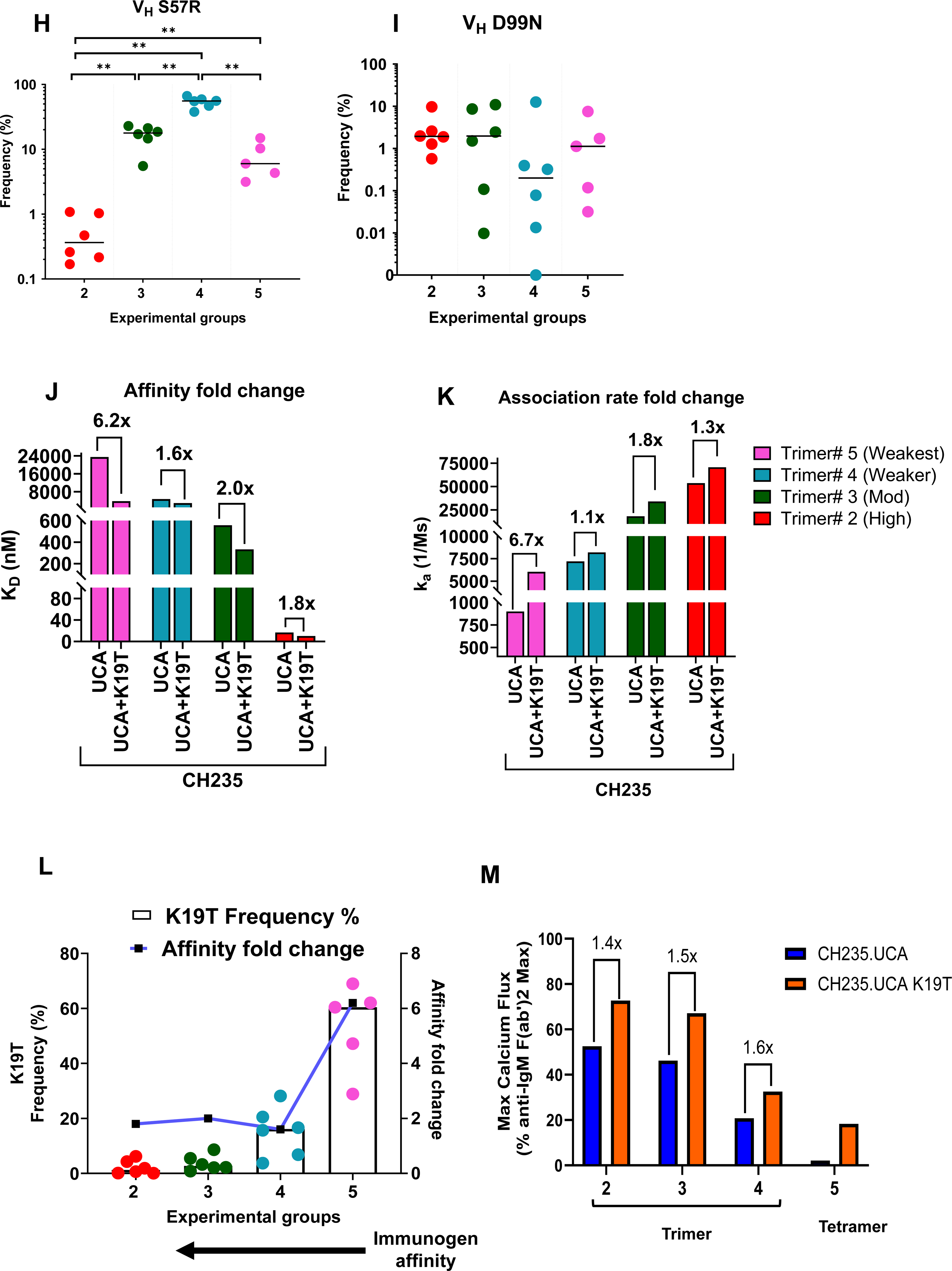
Relationship between affinity and frequency of key functional mutations. **(A)** Heavy chain amino acid mutations in the UCA to proximal intermediate I60 and early intermediate I59 in the CH235 lineage maturation pathway is annotated with the amino acid changes and colored according to their mutation probability in the CH235 bnAb clonal tree. Mutation probability <2% was classified as improbable by ARMADiLLO. Arrow indicates specific mutation and color coded based on mutation probability. **(B)** Variable heavy-chain (V_H_) median mutation frequency of immunized mouse knockin loci derived IgG sequences. **(C)** Mean number of heavy-chain improbable amino acid mutations in knockin loci derived IgG sequences in various experimental groups. **D-I** Plots show mutation frequencies (%) in experimental groups 2-5. Different V_H_ mutations observed in the CH235.UCA to I60 and I59 intermediates are shown as: **(D)** K19T (I60; improbable), **(E)** N52D (I60; less improbable), **(F)** S31N (I60; probable)**, (G)** M34I (I60; probable)**, (H)** S57R (I59; probable)**, (I)** D99N (I59; improbable). Note: for the experimental group#5, samples from five mice were sequenced. In each of the above plots the bar across each group represents the median value of the calculated frequency or number of improbable mutations. **(J)** Affinity (K_D_ ; nM) of trimers #2-5 to CH235.UCA and CH235.UCA+K19T mutated Fab. The plot shows fold change in affinity between CH235.UCA and UCA+K19T. **(K)** Association rates (k_a_ ; 1/Ms) of trimers #2-5 to CH235.UCA and CH235.UCA+K19T Fab. Fold change in k_a_ plotted between CH235.UCA and UCA+K19T. **(L)** Frequency (%) of the K19T mutation and affinity fold change in immunized groups 2-5. The plot demonstrates the K19T frequency (%) (y-axis1) versus affinity fold change (y-axis2) in immunized groups 2-5 (x-axis). Line connects the fold change values for each immunized group. **(M)** Maximum Ca-flux response for trimers #2, 3, and 4 in CH235.UCA (blue) and CH235.UCA+K19T (orange) Ramos cell lines. The fold change in the peak Ca-flux response between CH235.UCA versus CH235.UCA+K19T Ramos cell lines is indicated directly above the plotted responses. Calcium activation was observed with trimer #5 and the CH235.UCA+K19T Ramos cell line when the trimer was tetramerized via streptavidin and this peak Ca-flux response is shown in the left most bars. Statistical analysis was done using Mann-Whitney test. Statistically significant differences between groups are shown by asterisk (\**P*<=0.05, \*\**P*<=0.01). *P-*values for functional mutations are given in Tables S14 and S15.

First, we evaluated IgG sequence mutations, starting with summary metrics. The VH nucleotide mutation frequencies of all mice in the four groups are between 2.1 and 4.1 percent. Group 4 has a statistically significantly higher median mutation frequency than that of group 3 and 5 (group 4 > 3 p=0.0238; group 4 > 5 p=0.0043). Group 2 also has a statistically significantly higher median mutation frequency than that of group 5 (group 2 > 5 p=0.0173) (**Fig. 4B, Table S14**).

We also looked specifically at improbable amino acid mutations. The range of the mean number of improbable mutations for all the mice is between 3.45 and 4.44. Group 3 has a statistically significantly higher median value for the mean number of improbable mutations than group 2 (group 3 > 2 p=0.0087). No other group comparisons reached statistical significance (**Fig. 4C, Table S14**). The relatively small differences between these groups at the level of summary metrics suggested the presence of specific mutations could be responsible for the differences in neutralization.

One key improbable V_H_ mutation in the CH235 lineage is K19T which is inferred to occur very early in the development of the lineage in the UCA-proximal/early intermediate I60 (**Fig. 4A**). Inclusion of the V_H_ K19T substitution enhances affinity of CH235.UCA antibody and contributes to neutralization against autologous viruses^28^. The K19T mutation is also found in most VH1-46 CD4-bs class bnAbs underscoring its functional role in the recognition mode of this bnAb class^24,28^. Earlier studies showed that a UCA-targeting immunogen can select for K19T mutation^28^, and therefore, we first examined the frequency of the key improbable K19T mutation in each of the four immunized groups. We observed that the K19T mutation was not restricted by affinity, and the mutation was detected over the full range of immunogen affinities (µM – nM) in the panel, although with varying % frequency. An inverse linear relationship between affinity and K19T mutation frequency was observed, with the weaker affinity immunogens (groups 4-5) inducing significantly higher mutation frequency (group 4 > 2 p=0.0087; group 4 > 3 p=0.0152; group 5 > 2, 3, and 4 p=0.0043) (**Fig. 4D, Table S14**).

To address if additional mutations contributed to the observed differential neutralization in the immunized groups and whether there are affinity limits (lower/upper) that favor functional mutation selection, we analyzed the frequency of key mutations that were inferred in the UCA-first (I60) and second (I59) early intermediates of the CH235 lineage (**Fig. 4A**). The binding of CH235 bnAb to the CD4-bs epitope relies on contacts that are important for viral neutralization breadth and include the CD4-bs loop, the loop D and the conformationally variable loop 5^30^. In addition to K19T, the UCA-first intermediate I60 includes a key improbable (1-2%) V_H_ N52D mutation, and two probable mutations (2-10%), V_H_ S31N and V_H_ M34I (**Fig. 4A**). The mutational analysis in the different immunized groups showed that the frequency of V_H_ N52D was significantly higher in group 4 (median 3.15%) than either the higher affinity (0.14%-group 2) (group 4 > 2 p=0.002) or the weakest affinity group (0.02%-group 5) (group 4 > 5 p=0.0043) (**Fig. 4E, Table S14**). The median frequency of V_H_ N52D in group 3 was not dissimilar to that in group 4 (3.15% and 3.63% groups 4 and 3 respectively), but unlike that in group 4 (6/6 mice), the higher frequency was not observed in each of the immunized mice in group 3 (4/6 mice). The frequency of V_H_ S31N was not significantly different in the immunized groups (**Fig. 4F, Table S14**). However, the median frequency of the more probable V_H_ M34I was lower in groups 3 and 4 when compared to the higher and the weakest affinity groups (group 2 > 3 p=0.0411; group 5 > 3 p=0.0087; group 5 > 4 p=0.0303) though the differences in medians were modest (**Fig. 4G, Table S15**). Each group elicited both probable and improbable functional mutations, but with slightly different median mutation frequency range (S31N, M34I = 15%-37%; K19T, N52D = 0.1%-53%).

Next, we looked at mutations in the second intermediate I59 that includes a probable V_H_ S57R and an improbable V_H_ D99N (**Fig. 4A**), that play a role in adapting to flexibility in the variable V5 loop^30^. The highly probable V_H_ S57R mutation was observed in each of the immunized groups, but with the median frequency increasing with a decrease in affinity and then declining in the weakest affinity group (**Fig. 4H**). Both the moderate and weaker affinity groups 3 and 4 (0.56 and 4.7μM) showed significantly higher frequency than the groups with higher (16.7nM) (groups 3 and 4 > 2 p=0.0022) or the weakest affinity (23.5μM) (group 3 > 5 p=0.0519; group 4 > 5 p=0.0043) (**Fig. 4H, Table S15**). The highest median frequency of V_H_ S57R was observed in group 4 and that was significantly higher than in group 3 (group 4 > 3 p=0.0022). The improbable V_H_ D99N median frequencies were not significantly different between the immunized groups (**Fig. 4I, Table S15**). Additional mutations inferred in two early intermediates (I60 and I59) were low frequency except V_H_ S59N which had significantly higher frequency in group 2 compared to the rest of the groups (**Fig. S11A-E, Tables S15-S16**).

To explain the linear relationship between decreasing affinity and higher selection of key improbable V_H_ K19T mutation, we compared UCA Fab affinities and kinetic rates to each trimer protein to that of the mutated CH235.UCA_K19T Fab (**Fig. S12A-B**). The highest affinity gain (6.2-fold) with K19T mutation was observed with the weakest affinity trimer #5 (**Fig. 4J**) and this affinity gain was due to an equal increase in the magnitude (6.7-fold) in the binding on-rate (**Fig. 4K-L**). Although with a relatively lower K19T

frequency when compared to group 5 (60.4%), B cells in group 4 mice (16.18%) had a significantly higher K19T frequency than the two higher affinity groups (1.25% and 2.72% in groups 2 and 3 respectively). However, the affinity/on-rate gain was low in group 4 and similar in fold-increase to that in groups 2-3 (**Fig. 4J-K**). The more favorable selection of K19T in group 4 over the higher affinity groups 2-3 (**Fig. 4L**) could be explained if the BCR signaling that is below a threshold is more favorable for mutation selection, and/or that the selection of other functional mutations acts synergistically in driving more favorable selection. The former hypothesis is consistent with the observation that while peak Ca-flux responses in Ramos cells expressing BCRs with the K19T mutation (CH235.UCA+K19T) increased by 2-fold in response to each immunogen, the BCR signaling magnitude induced by the weaker affinity immunogen remained relatively lower (trimer #3 4 at <40%) than that with the higher affinity immunogens (trimer #1-2 2-3 at ∼70%). Additionally supporting this hypothesis, we observed that the weakest affinity immunogen when tetramerized induced weak, but observable Ca-Flux responses (tetramerized trimer #5 at ∼15-20%) with the CH235.UCA+K19T Ramos cells whereas no signaling was observed with the cells expressing BCRs of CH235.UCA (**Fig. 4M**). Overall, these results show that the selection of the key improbable K19T mutation is favored by the magnitude of gain in affinity due to an on-rate increase but biased towards weaker affinity (μM) immunogens and therefore, indicating the likelihood of an affinity/signaling ceiling for selection of functional mutations.

Finally, we asked if the frequency of combinations of functional mutations is different in the four immunization groups. We focused on the combination of three key mutations (K19T, N52D, S57R) that were each observed at relatively higher frequency in our immunization studies. The frequency of the K19T+N52D+S57R combination was significantly higher in group 4 when compared to either the higher affinity group 2 or the weakest affinity group 5, with most animals (5/6) in the two latter groups showing no combined mutations (group 4 > 2 p=0.0022; group 4 > 5 p=0.0390) (**Fig. S11F, Table S16**). While group 3 median was the same as group 4 frequency median, 50% of the group 3 animals showed either no mutations or much lower frequency, indicating a bias favoring higher frequency of the triple mutations in the weaker affinity group 4.

Taken together, the mutational frequency analysis showed that key functional mutations that were identified in the early intermediates of CH235 lineage were more favorably selected in response to immunogens that were in the mid-affinity range (0.5 - 5μM) but with a strong bias towards the weaker affinity group. While one functional mutation (K19T) was most favorably selected by the weakest affinity immunogen, selection of multiple key mutations together is not favored by either the high or the weakest affinity immunogens. Furthermore, the neutralization data is also consistent with this mutational analysis, and thus further supports the notion that BCR affinity/signaling imposes boundaries (both upper and lower limits) and that the selection of antibody functional mutations is most favorable in response to immunogen affinities within a narrow range.

In summary **(Fig. 5**), our results show that the strength of BCR signaling is proportional to antigen affinity/on-rate (**Fig. 5**, top panel). However, the viral neutralization titer increased with decreasing signaling strength but dropped precipitously in response to the immunogen with the weakest signaling, indicating that the pseudovirus neutralization is not dependent on the magnitude of sera binding titers and that there is a narrow BCR signaling range that favors induction of high neutralization titers (**Fig. 5**, top panel). The analysis of antigen-specific GC B cell frequency, and the frequency of key functional mutations gave results consistent with the preferred narrow range of signaling that induced higher neutralization (**Fig. 5**, middle panel). Our studies support the model (**Fig. 5**, bottom panel) that B cell signaling sets an upper and lower boundary for selection of functional mutations. Thus, B cells within a narrow range, the “Goldilocks zone” (0.5-5.0μM in the studied model and vaccine regimen), affinity/signaling are favored for longer GC dwell time and LZ-DZ cycling, and SHM driven selection of key functional mutations, and subsequently in the induction of antibody responses with high neutralization potency. Conversely, too strong BCR signaling is disfavored for selection of functional mutations due to limited LZ-DZ cycling and an early GC exit into the plasmacytic compartment (**Fig. 5**, bottom panel).

**Figure 5.**
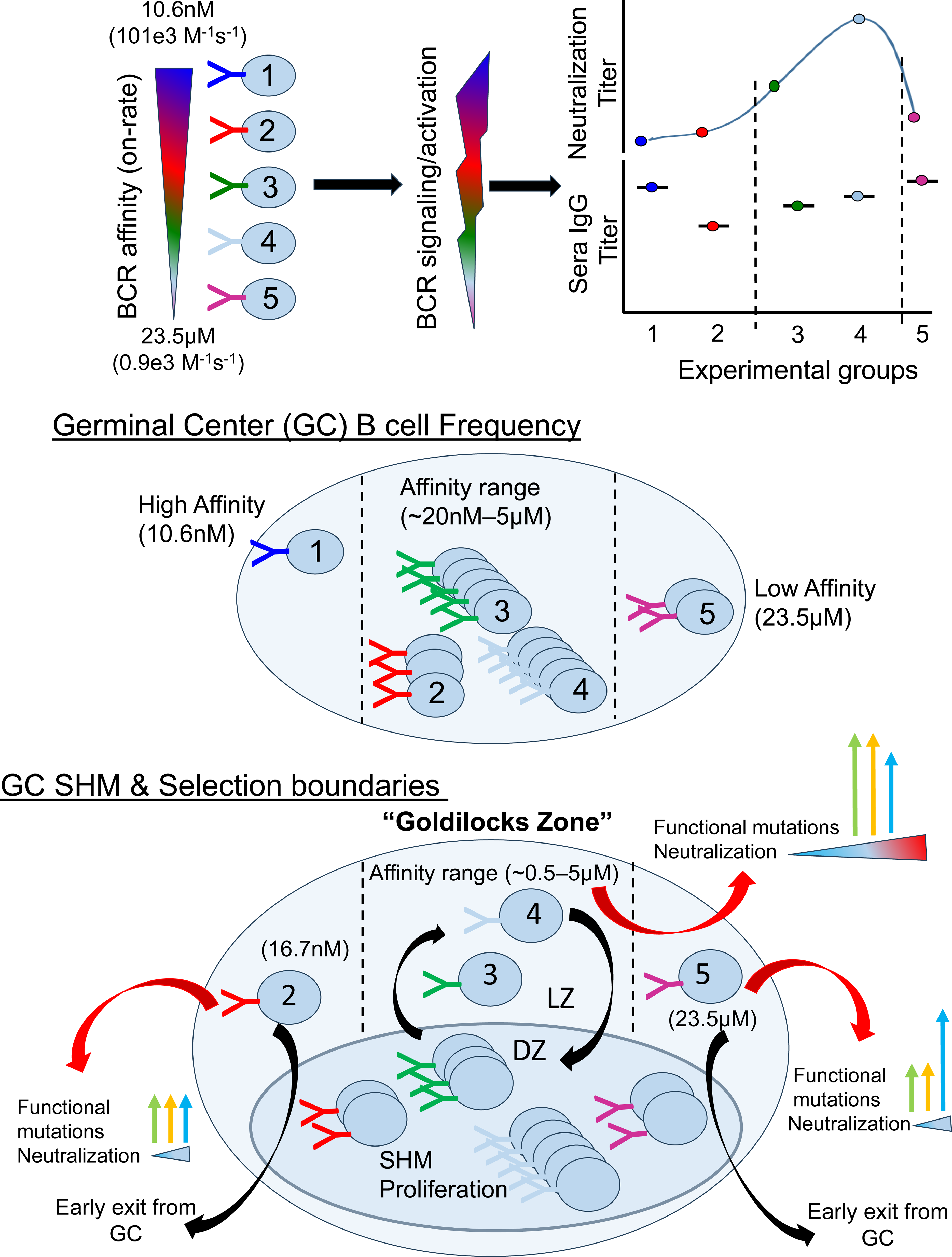
The “Goldilocks Zone” model for the selection of antibody functional mutations and neutralization. **Top panel:** B cell receptor (BCR) signaling strength increases with increasing trimer affinities (K_D_; nM-µM) primarily due to an increase in binding association rates (on-rate) (k_a_; M^-1^s^-1^). Sera neutralization titers are higher within the moderate (0.56-4.7 μM) affinity group and favors the weaker affinity group indicating that the neutralization titers are not dependent on the magnitude of sera IgG antibody titer. **Middle panel:** Higher GC B cell frequency within a narrow affinity range (∼20nM-5µM) and biased against both the highest and the weakest affinity groups. **Bottom panel:** For key antibody functional mutation selection, the affinity range gets narrower with B cells within the “Goldilocks zone” (∼0.5-5µM) being the most favored group. B cells with affinities outside the favored zone (both higher and lower) acquire fewer mutations, likely due to shorter GC dwell time and LZ-DZ cycling, consequently, induce antibodies with weak neutralization titers and exit earlier from GC into the plasmacytic compartment. The height of the arrows represent percent mutation frequencies (green-N52D, orange-S57R, blue-K19T). A color gradient represents the neutralization titer. Trimer #1 (Highest affinity; 10.6nM; dark blue), Trimer #2 (High affinity; 16.7nM; red), Trimer #3 (Moderate affinity; 557.3nM; green), Trimer #4 (Weaker affinity; 4700nM; light blue), and Trimer #5 (Weakest affinity; 23.5µM; pink).

## Discussion

Development of a protective anti-viral humoral response is dependent on the AID-mediated SHM process that drives the occurrence of mutations in the germinal center of B cell follicles^32^. The selection of functional mutations is critical for the development of broadly neutralizing antibodies that target the conserved sites on the HIV-1 Envelope trimer protein, including the CD4-bs epitopes^28^. Following immunization studies in a CD4-bs bnAb UCA-specific humanized mouse model, we made the central observation that B cell signaling strength sets boundaries for both the selection of key functional mutations and sera viral neutralization potency. Our studies support the “Goldilocks zone” model (**Fig. 5**) and define the BCR signaling strength that is most favored for selection of antibody functional mutations and viral neutralization.

In our earlier studies, we had reported that BCR signaling strength is dependent on antigen binding association rate (on-rate)^20^ and here we further validate it in a distinct CD4-bs bnAb UCA model. Since association rate increases are observed in early bnAb maturation^33,34^, an on-rate dependent affinity gain will favor positive selection of a mutation^30^. A key finding here is that an increase in association rate and not the starting affinity gave the highest frequency of a critical mutation (K19T) that enhances autologous neutralization, a finding highly relevant in designing priming HIV-1 immunogens^30^. Our results showing the gain in affinity as the driving factor in mutation selection implies that GC B cells are subject to relative affinity comparison^13,14,35^ and a lower starting affinity will provide more opportunities for higher gain, and consequently providing more of a selective advantage to B cells harboring the specific mutation which will lead to higher frequencies of that mutation. However, our studies show that while the weakest affinity group 5 has the highest frequency of one critical mutation (K19T), this group was inferior for other key single or combined mutations (K19T, N52D, S57R), and as a result induced weak neutralization titers. The above result suggests that B cells with a limited number of mutations (as in group 5) likely reach a roadblock in affinity maturation either due to exclusion by competition with continuously entering (off-target, comparably low affinity) naive B cells into existing GCs^36^, or because they lack the intrinsic signaling capacity to activate their cellular B-cell GC proliferative and GC differentiation programs^37^ and as a result the average affinity in response to group 5 immunogen remained lower (**Fig. 2, S6**).

The role of BCR signaling in mutation selection is dependent on the endocytic function of the BCR that provides GC B cells with the ability to receive positive signals from TFH cells. In a recent study, it was shown that BCR signaling per se is required for LZ B cell survival and positive selection leading to DZ migration and proliferation^10^. The range of affinity that triggers BCR signaling spans from μM to low nM (10^-6^ to 10^-10^ M), and when not limited by low precursor frequency or competition with higher affinity B cells, affinity as low as 100μM can be sufficient to activate naïve B cells and initiate GC responses^13,15^. However, the range of affinities that is permissible for the selection of functional mutations has not been clearly defined. In this study, our first observation that B cell signaling differentially modulates GC B cell responses came from the phenotypic analysis of B cells showing that the frequency of class-switched and trimer-specific IgG+ GC B cells increased with decreasing trimer affinity till the lower threshold was reached with the weakest affinity group 5. The above trend was mirrored in the results from the analysis of TFH cells (CXCR5+PD1+) and therefore indicated that GC B cell signaling bridges B-T cell responses^10^. The impact of starting affinity was evident in the inverse relationship between BCR signaling and the selection of a critical mutation, K19T. Our results clearly show that too high an affinity is disfavored and that is consistent with the differentiation outcome favoring early GC exit into the extracellular plasma cell (PC) compartment^18,19^. At the low affinity end and with respect to functional mutations and neutralization, we observed a similar outcome with the weakest affinity group 5, although as discussed above likely due to a different mechanism in play. These results support our hypothesis that there is an upper/lower limit of BCR signaling in selection of antibody functional mutations that are required for viral neutralization (**Fig. 5**). The frequency of several key mutations (K19T, N52D, S57R), both as single or a combined set of mutations, was observed to be highest in the mid-affinity group and lowest in groups at either end of the affinity spectrum. The differences in the antibody mutation frequency in the immunized groups were consistent with the observed bell-curve profile of the neutralization potency, validating that moderate BCR signaling favors selection of key functional mutations. The observed low frequency of mutations in the high affinity disfavored group (group 2) could be due to minimal LZ-DZ cycling, shorter LZ residency and earlier exit to the PC compartment (**Fig. 5**), and consistent with the model that PC differentiation is restricted to the highest affinity B cells^19,38,39^. The low levels of functional mutations in both group 2 (high affinity) and group 5 (weakest affinity) indicate that B cells in both groups have shorter GC dwell times^36,40^. Overall, we have defined the range of BCR affinity/signaling that is most favored for selection of functional mutations, and our results support the GC fate decision model wherein intermediate affinity B cells (The “Goldilocks Zone”) reenter the DZ and proliferate, while the high affinity exit earlier to the PB/PC compartments^1,11^.

One potential limitation in our study is the use of a humanized mouse model that populates relatively higher frequency (∼5-10% of follicular B cells)^28^ of Env trimer-binding mature B cells and presents a caveat that the defined “Goldilocks zone” might need to be redefined in the setting of lower precursor frequency. Studies on the role of antigen affinity/valency in B cell responses and differentiation report the requirement of relatively higher affinity (0.5μM) and a high-order valency form (60-mer) when the targeted B cell precursor frequency is low^16,41^. However, a recent study has shown a lower affinity eOD-variant of the eOD-GT platform, eOD-GT7 (∼3 μM) outperforms the much higher affinity eODG-GT8 clinical candidate (∼15 nM) in overall GC fitness and improbable SHM acquisition across all frequencies tested (including comparable frequencies of bnAb precursors this model expresses, all the way to ultra-low ie, ∼1/10^6^ reconstitutions) after the prime step in a bnAb precursor adoptive transfer model^37^, suggesting this feature is at least partly intrinsic to the antigen-BCR sensing mechanism itself, irrespective of low affinity/off-target clonal competition. Regarding valency specifically, a much lower affinity (40µM) protein in a low valency form (trimer) has been reported to be effective with as low as 10 knock-in precursor B cells in terms of knock-in B cells recruitment into the GC, though B cell expansion was limited and recruitment was more variable compared to the high affinity trimer (7nM)^42^. It will be intriguing to further test valency requirements across multiple bnAb lineage targets and with different immunogen forms (trimers, scaffolds, etc.) as differing valency requirements might be influenced by not only differences in baseline BCR signaling thresholds of peripheral B-cells populations imparted by prior encounter with self-antigens^43,44^, but the maturational stage of bnAb development the knocked in V(D)J bnAb rearrangements represent ^16,42^ and/or the minimal protein form used as the immunogen.

Finally, this study was specifically designed to focus on the long-term functional outcomes of serum neutralization and improbable somatic mutations from sampled memory IgG+ B-cells, after several rounds of recall and selection (from immunogens being compared in this study used a full, four shot vaccine regimen). While we show increased terminally-sampled antigen-specific GC+ B-cell compartments correlate with neutralization and SHM improvements in response to immunogens that fall within the ‘goldilocks zone”, it will be intriguing in follow-up these studies with those examining the relative impact of initial recruitment/cycling of bnAb-specific B-cells in GCs (during priming) relative to those re-entering GCs during subsequent recalls. In this regard, recent complementary work in a chimeric CH31 UCA transfer model demonstrate a large effect on an analogous affinity floor/ceiling stemming from GC-specific transcriptional programming as early as 24 hours ex vivo and predictive of peak GC fitness to in vivo priming^37^, suggesting the affinity gains observed here from CH235 BCR sensing of lower affinity antigens may not only be due to off-target follicular competition and/or GC LZ selection, but more fundamentally, due to directly modulating the crucial processes of survival and SHM in DZs, suggesting in turn that GC fate programming have the potential to eventually be predictively tuned based on pre-defined characteristics of any antigen-BCR pair’s defined biophysical binding kinetic properties.

## Author Contributions

S.M.A. conceptualized and designed experiments. S.M.A., A.S. and K.A. prepared the manuscript with input from K.W., E.V.I. and L.V. and edits from all authors. S.M.A. supervised research. K.A. performed SPR binding assays, immunogen QC, and calcium flux measurements. A.S., A.N. and the DHVI immunization team conducted the immunization studies, sera and tissue harvesting. A.S. performed ELISA, immunophenotying, ex-vivo calcium flux assay, analyzed binding titer data and flow cytometric analysis. A.P.K. performed Ramos cell line QC and phosho-flow kinase assay. M.B. and R.P. performed ELISA and analyzed data. K.W., S.V., E.V.I. analyzed NGS data and mutation frequency analysis. D.W.C. performed flow cytometric cell phenotypic analysis. R.H. and K.S. designed and produced HIV Envelope proteins. F.W.A. and M.T. developed the CH235.UCA knock-in mice. S.M.A., L.V., B.F.H. provided resources and funding.

## Acknowledgements

We thank Dr. Bart Haynes, Director at DHVI for providing facility resources at respective sites and for scientific advice. We thank the DHVI protein expression facility (Director: Kevin Saunders), the DHVI flow cytometry and sequencing facilities, and the DHVI finance and administrative teams, for their support. We thank Darrel Irvine, Scripps Research Institute, La Jolla, CA, for providing the SMNP adjuvant. We thank Katrina Hodges and Chuancang Jiang for neutralization assay and Letealia Oliver for immunophenotyping. We also acknowledge Wes Rountree for performing statistical analysis.

Research reported in this submission was supported by the National Institute of Allergy and Infectious Diseases (NIAID) of the National Institutes of Health (NIH) under Award Numbers R01 AI145656 (SMA). BFH was supported by the NIAID Consortia for HIV/AIDS Vaccine Development grant UM1 UM1AI144371. The content is solely the responsibility of the authors and does not necessarily represent the official views of the National Institutes of Health.

## Methods

### Study design

The major objective of this study is to evaluate the impact of varying affinities and kinetic rates on HIV-1 envelope (Env) proteins immunogenicity with respect to selection of functional mutations and autologous viral neutralization) in knock-in (KI) murine model. The CD4 binding site (CD4-bs) of the HIV-1 Env proteins is target for multiple lineages of broadly neutralizing antibodies (bnAbs). The CH235 (Vh1-46 CD4-bs) lineage of CD4-bs bnAbs is of particular interest as it evolved to 90% neutralization breadth, with no indels and normal CDR3 lengths^25,26^. In addition, CH235.UCA binds with higher affinity to CH505.TF trimers with selected mutations (N279K, G458Y) and can neutralize such mutant pseudoviruses^27^. Moreover, critical improbable mutations acquired early in CH235 development have been identified^28^. To this end, we selected CH505.M5/TF Env trimer immunogens and chose CH235.UCA KI mice as the model to test immunogenicity, as it can particularly select functional somatic mutations required for neutralization. We hypothesized that a prime immunogen with the optimal affinity/rates and signaling strength will favor selection of functional mutations and induction of autologous neutralization. Here, in sequential homologous immunization studies in CH235.UCA KI mice, we used 5 Env trimers that bind with varying affinities/rates and investigated the role of B cell signaling strength in priming selection of functional mutations and autologous pseudovirus neutralization.

### Antigens

The following HIV Env gp140 trimer proteins were produced at the Duke Human Vaccine institute (DHVI) as described previously and includes the following:

CH505M5chim.6R.SOSIP.664v4.1_G458Y_D403C+T411C_N332R_N197D_F14_Wpro (trimer #1)^30^, CH505M5chim.6R.SOSIP.664v4.1_G458Y_D403C+T411C 293F (trimer #2)^30^, CH505M5chim.6R.SOSIP.664v4.1_G458Y 293F (trimer #3)^27^, CH505M5chim.6R.SOSIP.664v4.1 (trimer #4)^27^, and CH505TFchim.6R.SOSIP.664.v4.1_G458Y (trimer #5)^29^. Three CH505 trimers (Trimer #1, 2 and 3) in the antigen panel were engineered with both the N279K (M5 variant) and G458Y mutations that enhance the affinity of the CH235.UCA antibody and its ability to neutralize^27,28^, while trimer #4 had only the M5 mutation and the weakest affinity trimer #5 had G458Y but not the M5 mutation.

### Surface Plasmon Resonance (SPR) Affinity and Kinetic Rate Measurements

The SPR affinity measurements (K_D_) and kinetic rates (k_a_ and k_d_) for CH505 Env trimers were obtained using the Biacore S200 or T200 instrument (Cytva) in HBS-EP+ 1X running buffer as previously described^27,28^. Biotinylated trimer proteins were immobilized to a level of 300-500 RU onto an SA (streptavidin) sensor chip (Cytiva) while non-biotinylated trimers were captured onto PGT151 immobilized (5000-10000RU) chip surfaces to a level of 200-1000RU. CH235 lineage Fabs were used as the analytes for all trimers. CH235.UCA and other lineage Fabs (I60, I59, and I39) as well as CH235.UCA_K19T Fab were diluted from 50nM to 3000nM and injected over the sensor chip surfaces using the single cycle injection type. Five sequential injections of the CH235 lineage Fabs were injected at a flow rate of 50uL/min for 120s per injection and were followed by a 600s dissociation period. Regeneration with a short pulse of Glycine pH2.0 was used for all trimers. Binding to a blank streptavidin or CM5 chip surface as well as buffer binding were used for double reference subtraction, non-specific binding, and signal drift. A 1:1 Langmuir model with a local Rmax was used for the Fab curve fitting analysis against most trimer proteins. The heterogeneous ligand model was used for CH235.UCA Fab binding to HV1302590 with the faster kinetic rates reported. The reported kinetic rate and affinity values are representative of 2 data sets.

### Cell lines

Ramos cell lines were developed as described previously with slight modifications^45,46^. CH235.UCA Ramos B cells were developed with the use of lentiviral transduction. Plasmids containing the CH235.UCA variable region and IgM membrane bound DNA sequences were synthesized using PCR and bacterial preps (Wizard Plus SV Miniprep DNA Purification System, Promega). These plasmids were then transfected into 293T cells for lentiviral production. After stable transduction of naive Ramos cells, cells were stained with fluorescent antibodies targeting the Fcɣ and ĸ light chain portions of the BCR and phenotyped using BD LSRFortessa flow cytometer. The cells were then enriched two times to select for the highest double positive expressers of BCR. A final phenotype was done to ensure expression was at the same level as other BCR expressing Ramos cell lines. If expression was lower, then another sort was conducted.

### Calcium Flux in Ramos B cells

Calcium flux experiments were performed using the FlexStation 3 Microplate Reader (Molecular Devices) in conjunction with the FLIPR Calcium 6 dye (Molecular Devices) as previously described^46^. A cell count for the Ramos cells was performed with a Guava Muse Cell Analyzer (Luminex) to ensure cell viability was greater than 95% and to calculate the volume of cells needed for a concentration of 1x10^6^cells/mL on the day of the experiments. The required volume of cells was then pelleted at 1500rpm for 5 minutes. The supernatant was then decanted and the cells were resuspended at a 2:1 ratio of RPMI media (gibco) + FLIPR Calcium 6 dye (Molecular Devices). The cells were plated in a clear, U-bottom 96-well tissue culture plate (Costar) and incubated at 37°C, 5% CO_2_ for 2h. Antigens were separately diluted down to a concentration of either 200nM or 500nM in 50uL of the 2:1 ratio of RPMI media (gibco) + FLIPR Calcium 6 dye (Molecular Devices) and plated in a black, clear bottom 96-well plate. The final concentration of antigen would be 100nM or 250nM based on the additional 50uL of cells added during the assay. A positive control stimulant, Anti-human IgM F(ab’)_2_ (Jackson Immuno, cat 109-006-129) diluted down to 100ug/mL (50ug/mL final concentration) was also included in the antigen plate. Using the FlexStation 3 multi-mode microplate reader (Molecular Devices), 50uL of the cells were added to 50uL of protein or Anti-human IgM F(ab’)_2_ diluted in RPMI/dye and read continuously for 5min. The fluorescence of a blank well containing only the RPMI/dye mixture was used for background subtraction. After subtraction the antigen fluorescence was then normalized with respect to the maximum signal of the IgM or IgG control and calcium flux values were presented as a percentage. Calcium flux data are representative of at least 3 measurements.

### In-vitro Phospho-flow kinase assay

Phosphorylation of three proximal kinases (pSyk, pBtk, pBLNK) were analyzed in Ramos cell line as described previously with slight modifications^20^. Briefly, a total of 2.5x10^6^ CH235.UCA IgM Ramos cells were stimulated with each trimer at 100nM at various time points over a 30-minute period in a thermomixer (Eppendorf) at 37°C. Cells were collected after each period of incubation followed by fixation, permeabilization and staining with fluor conjugated anti-human Syk (BioLegend, cat 683708), Btk (BD Biosciences, cat

564848), and BLNK (BD Biosciences, cat 558444) antibodies. Cells were washed and resuspended in PBS/1%BSA (0.05% NaN3) buffer. Data was acquired using BD LSRFortessa flow cytometer and analyzed using FlowJo software (v.10.8.1). Normalized MFI for each timepoints were obtained dividing treated sample MFI by unstimulated or 0 min sample MFI.

### CH235.UCA knock-in mice

The humanized murine model was developed with the V_H_DJ_H_ and V_λ_J_λ_ regions of the CH235.UCA knocked in and were maintained on C57BL/6J backgrounds as heterozygous lines as described previously^28^. All mice were placed into standard housing system at DHVI animal facility in accordance with NIH guidelines. The experimental procedures were approved by Institutional Animal Care and Use Committee (IACUC).

### Immunization

A total of 31 eight to twelve-week-old CH235.UCA V_H_/V_L_ mice (heterozygous) were randomly divided into two studies. Study #1 and #2 were divided into three and two experimental groups, respectively (n=6 per group). Mice in both the studies received 25ug of CH505 trimer proteins intramuscularly four times at two-week interval apart, formulated in 5ug of saponin/monophosphoryl lipid A nanoparticles (SMNP) adjuvant. Study #1: group #2-4 received trimer #2-4, respectively. Study #2: group #1 and #5 received trimer #1 and #5, respectively. Both wild-type C57BL/6 and knock-in CH235.UCA were kept as control mice (group #0) and received 1X PBS. Blood samples were collected at one-week post each immunization and samples were used to evaluate IgG antibody binding titer by enzyme-linked immunosorbent assay (ELISA). At day 7 post last immunization mice belonging to each group were euthanized and tissues spleen and lymph nodes were harvested. Sera samples processed post fourth immunization were heat inactivated at 56 °C for 15 min for pseudovirus neutralization assay whereas spleen and lymph nodes were processed for the NGS analysis of functional mutations and immunophenotyping, respectively. Trimers used in this study are given in Table S1.

### Serum Enzyme-linked immunosorbent assay (ELISA)

The presence of antigen specific total IgG antibody level in the sera samples were analyzed by ELISA as described previously with minor modifications^47^. Briefly, 384 well-plates were coated with PGT151 antibody (2ug/mL) in 0.1M sodium bicarbonate buffer overnight at 4°C. The plates were washed with PBS-T (1X PBS with 0.1% Tween 20) followed by blocking (5mL 10XPBS, 7.5mL goat serum, 0.25mL tween 20, 0.5mL sodium azide, 2g whey protein and bring the volume upto 50mL using deionized water) for 1h at room temperature (RT). The plates were further treated with CH505.M5/TF trimer of varying affinities (2ug/mL) in blocking buffer and incubated for 1h at RT. Post incubation, serially diluted sera samples (starting with 1:30 dilution; followed by 3-fold dilutions) were added to wells and incubated for another 90min at RT. The plates were washed twice and incubated with HRP-conjugated goat anti-mouse IgG secondary antibody (1:10,000 dilution in blocking buffer without sodium azide) for 1h at RT. After washing four times the plates were developed with 3, 3′, 5, 5′-tetramethylbenzidine (TMB) substrate for 15min. The reaction was stopped with 0.33N HCl and absorbance was read at 450 nm using microplate reader. Binding titers are presented as log area under the curve (log AUC). Log AUC was measured based on the OD_450_ values of the starting dilution till the endpoint titer that was calculated based on the cut-off (average of blank x 3).

### Lymph nodes B and T cell phenotypic analysis by flow cytometry

To determine the phenotypes and frequencies of Env-specific B cells and Tfh cells and examine B cell differentiation into GC, memory, and plasmablast (PB) compartments, inguinal and axillary lymph nodes were harvested from immunized mice and processed into single-cell suspensions. For B cells, a total of 2x10^6^ lymph node cells were resuspended into 100μL 1X PBS (with 2%FBS). The master mix of mAbs (BB700 anti-mouse IgG1,2,3 (BD Biosciences, cat 742115, cat 745969, cat 745825), PE anti-mouse GL7 (BD Biosciences, cat 561530), PE-CF594 anti-mouse CD93 (BD Biosciences, cat 563805), PE-Cy5 anti-mouse TER119 (BioLegend, cat 116210), PE-Cy7 anti-mouse IgM (BD Biosciences, cat 552867), APC-R700 anti-mouse CD19 (BD Biosciences, cat 565473), BV421 anti-mouse CD5 (BioLegend, cat 100629), BV510 anti-mouse IgD (BD Biosciences, cat 563110), BV570 anti-mouse CD11b (BioLegend, cat 101233), BV605 anti-mouse CD95 (BD Biosciences, cat 740367), BV650 anti-mouse B220 (BD Biosciences, cat 563893), BV711 anti-mouse CD138 (BD Biosciences, cat 563193), BV786 anti-mouse CD23 (BD Biosciences, cat 563988)) and antigen specific hooks (VB515/AF647-CH505.M5.G458YGnTI-gp120) were prepared at 2X concentration in 1X PBS (with 2%FBS). Next, 100μL of this 2X master mix was added to cells. Cells were incubated at 4°C for 20min followed by washing with 1X PBS at 1500rpm for 5min. Next, cells were resuspended into 100μL 1X PBS containing Near-IR Live/Dead dye (ThermoFisher) at 1:1000 dilution, and incubated for 20min at RT. After incubation cells were washed with 1X PBS (with 2%FBS). Similarly, Tfh cells were analyzed except 1x10^6^ cells were used to stain with the following fluor conjugated antibodies: FITC anti-mouse CD4 (BD Biosciences, cat 553047), BB700 anti-mouse CD69 (BD Biosciences, cat 566500), Biotin anti-mouse CXCR5 (BioLegend, cat 551960), PE-CF594 anti-mouse CD279 (BD Biosciences, cat 562523), PE-Cy5 anti-mouse TER119 (BioLegend, cat 116210), PE-Cy7 anti-mouse CD62L (BD Biosciences, cat 560516) , AF647 anti-mouse CD25 (BD Biosciences, cat 563598), APC-R700 anti-mouse CD8 (BD Biosciences, cat 564983), BV421 anti-mouse CD127 (BD Biosciences, cat 566300), BV510 anti-mouse CD3 (BD Biosciences, cat 563024), BV570 anti-mouse CD11b (BioLegend, cat 101233), BV605 anti-mouse Thy1.2 (BD Biosciences, cat 563008), BV650 anti-mouse NK1.1 (BioLegend, cat 108736), BV711 anti-mouse CD44 (BD Biosciences, cat 563971), BV786 anti-mouse B220 (BD Biosciences, cat 563894). Both B and T cells were fixed in 1X PBS/2% formaldehyde and stored at 4°C. The following day data was collected on a BD LSRFortessa flow cytometer and analyzed using FlowJo software. B cells (CD19+ B220+) with phenotypes GL7+/- CD138+, GL7+ CD138-, and GL7- CD138- were identified as early/late PB, GC, and memory B cells respectively^48^.

The formula used to calculate the relative proportion of GC B cells to PB either as a percent of total B cells or IgG+ Env sp. B cells is as follows:

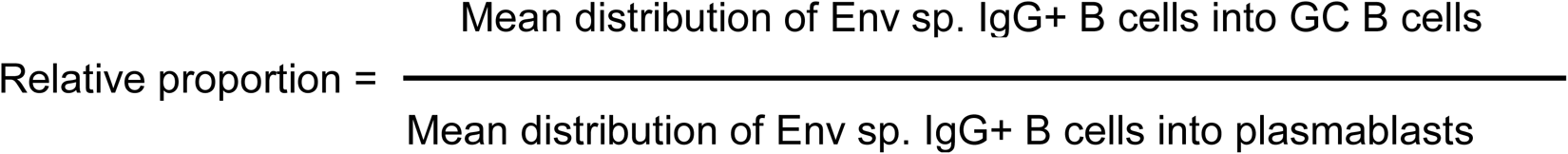

### Ex-vivo calcium flux analysis

The *ex-vivo* calcium flux assay for primary splenic B cells was performed as described previously with minor modifications^49^. Briefly, spleens from eight to twelve-week-old CH235.UCA KI mice were harvested. Single-cell suspensions from spleen were generated and incubated with 2mL ACK lysing buffer (Life Technologies) for 4min on ice to lyse red blood cells (RBCs). The splenocytes were resuspended (1x10^8^/mL) in HBSS buffer (gibco, cat14025-092) with 3% FBS. A total of 5x10^6^ cells were stained with BV650 anti-mouse B220 (1:100 dilution) (BD Biosciences, cat 563893) and APC-R700 anti-mouse CD19 (1:300 dilution) (BD Biosciences, cat 565473) and incubated for 40min at RT, protected from light. After washing, cells were resuspended in HBSS (5x10^7^/mL) and loaded with 2X Fluo-4 (Fluo-4 DirectTM Calcium Assay Kits, Invitrogen, cat F10471) in equal volume. The Fluo-4 mixed cells were incubated in bead bath at 37°C for 30min followed by incubation at RT for additional 30min. Cells were washed twice with HBSS (1mL/wash) and resuspended in HBSS (1x10^8^/mL). Further, cells were incubated with LIVE/DEAD Near IR dye (1:1000 dilution in DPBS) (gibco, cat 14190-136) for 30min at RT. Finally, cells were resuspended in calcium containing HBSS and stimulated with F(ab’)2-Goat anti-mouse IgM (MU chain) secondary antibody as a positive control (50ug/mL) (Invitrogen, cat 16-5092-85) and CH505 trimers 1-5 (50-100ug/mL) of varying affinities. Trimer CH505M5chim.6R.SOSIP.664v4.1_G458Y_N280D_n-avi-Bio/GnTI-(25ug/mL) was used as a negative control. Fluo-4 data for total B cells (CD19^+^B220^+^) was acquired using BD LSR II flow cytometer and analyzed by FlowJo software.

### Serum neutralization assay

Neutralizing antibody assays were performed in 96-well culture plates using HIV-1 Env-pseudotyped viruses and TZM-bl cells as previously described^29^. Serum samples were heat-inactivated at 56 °C for 15 minutes and initially diluted 1:10 in cell culture medium. Eleven microliters of the diluted serum samples were added in duplicate to the 96-well plates, followed by 3-fold serial dilutions. Env-pseudotyped viruses were produced by transfection in 293T cells. The samples were then incubated with virus for 1 hour at 37 °C in 5% CO₂ (resulting in a starting dilution of 1:200 for the serum samples). Subsequently, TZM-bl cells were added to each well and incubated for 48 hours. Following incubation, cells were lysed using Bright-Glo™ Luciferase reagent, and luciferase activity was measured with a microtiter plate luminometer. Neutralization titers were defined as the reciprocal of the serum dilution required to reduce relative luminescence units (RLU) by 50% compared to RLU in virus control wells.

### Next-generation sequencing (NGS)

Bulk single chain BCR repertoire sequencing of mouse splenocytes was performed as previously described^50^. Briefly, total RNA was extracted from mouse splenocytes using a RNeasy Plus Mini kit (Qiagen cat 74104), by automation on QIAcube instrument according to manufacturer’s protocol. RNA integrity and concentration was assessed using a Qubit Fluorometer (ThermoFisher). 1ug of total RNA was subjected to an Illumina compatible library prep using the SMARTer® Mouse BCR IgG H/K/L Profiling Kit (Takara cat 634422) using SMART technology, which employs a 5’ RACE-like approach to capture complete V(D)J variable regions of BCR transcripts. Briefly, the RNA was reverse transcribed with Poly dT provided in the SMARTer Mouse BCR kit for cDNA synthesis. Heavy and light chain genes were separately amplified using reverse primers that anneal in the mouse IgG constant region for heavy chain genes and IgK for the light chain genes. 5 µl of cDNA was used for heavy and light chain gene amplification via two rounds of PCR (18 and 12 cycles per round). During the second round of PCR, Illumina adapters and indexes were added. Illumina ready sequencing libraries were then purified and size-selected by AMPure XP (Beckman Coulter, cat A63881) using kit recommendations. The heavy and light chain libraries per mouse were indexed separately, thus allowing us to deconvolute the mouse-specific sequences during analysis. Libraries were quantified using QuBit Fluorometer (Thermo Fisher) and validated on a TapeStation 2200 (Agilent) using sing a D1000 HS kit (cat 5067-5584). Libraries were pooled by mice groups for sequencing on the Illumina MiSeq Reagent Kit v3 (600 cycle) (Illumina, cat MS-102-3003) using read lengths of 301/301. 20% PhiX was spiked in to increase sequence diversity of the libraries.

For experimental group #5, NGS was performed with a different protocol where total RNA was extracted from mouse splenocytes using the Qiagen RNeasy Mini Kit according to the manufacturer’s protocol. Libraries were prepared with the SMART-Seq Mouse BCR with UMIs kit (Takara Bio) according to the manufacturer’s protocol. Sequencing was performed on an Illumina NextSeq 2000 using a P2 600-cycle XLEAP sequencing kit.

### Processing of NGS sequence data

For libraries prepared with the Takara SMARTer Mouse BCR IgG H/K/L Profiling Kit, the processing steps were as follows: Paired NGS reads were merged, filtered so that only sequences having >=95% of bases with a Q score above 30 were retained, and deduplicated. Sequences were annotated with the Cloanalyst Program^51^. Sequences were assigned as knock-in when their V gene mutation frequency after assignment to a human V segment was lower than after assignment to a mouse V segment. Only functional (both invariant cysteines are present, invariant phenylalanine/tryptophan is present, indels must be modulo 3, and CDR3 is modulo 3) and productive (no stop codons) sequences were retained for downstream analysis. Mutation frequencies are reported at the nucleotide level for the portion of the V(D)J sequence up to and including the cysteine at the beginning of the CDR3.

For libraries prepared with the Takara SMART-Seq Mouse BCR (with UMIs) Profiling Kit, the analysis was the same as above except for the following: (1) sequences with the same UMI and <=5 bp of length variation were aligned and then corrected to the majority base (A,C,G,T,N) for each position or N if there was no majority base; (2) aside from grouping by UMI, there was no sequence deduplication; (3) sequences were assigned to an isotype (IgA, IgD, IgE, IgG, IgM, or other) based on alignment of the sequence 3’ of the VDJ to a library of the mouse heavy constant regions and only IgG assigned sequences were considered for further analysis.

### Statistical analysis

The GraphPad Prism version 10.1.2 was used to plot the graphs and for data analysis. Data for ELISA, phenotyping, and neutralization assay were compared across the immunized groups using exact Wilcoxon test with FDR correction. The median mutation frequencies induced by CH505 trimers of varying affinities were compared using Mann-Whitney test. Flow cytometric gating and data analysis for ex-vivo calcium flux and phenotyping were done using FlowJo software version 10.10.0. Data were considered statistically significant at \**P* < 0.05, \*\**P* < 0.01, \*\*\**P* < 0.001 (exact Wilcoxon test) whereas \**P* ≤ 0.05, \*\**P* ≤ 0.01, \*\*\**P* ≤ 0.001 using Mann-Whitney test.

