## Supplemental Figures for "B Cell Receptor Signaling Sets Boundaries for Selection of Broadly Neutralizing Antibody Functional Mutations"

### Slide 1
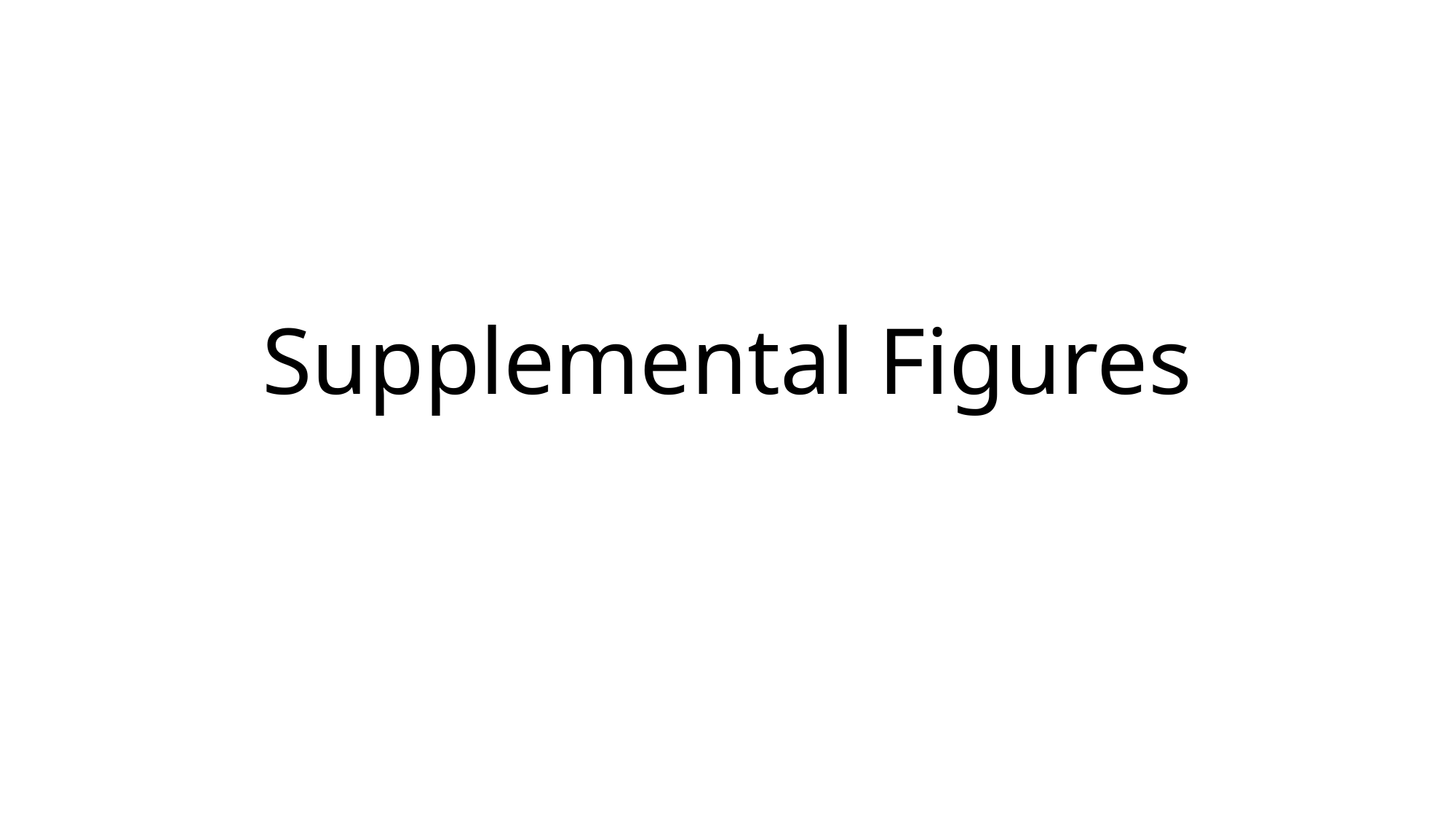

#### Supplemental Figures

### Slide 2
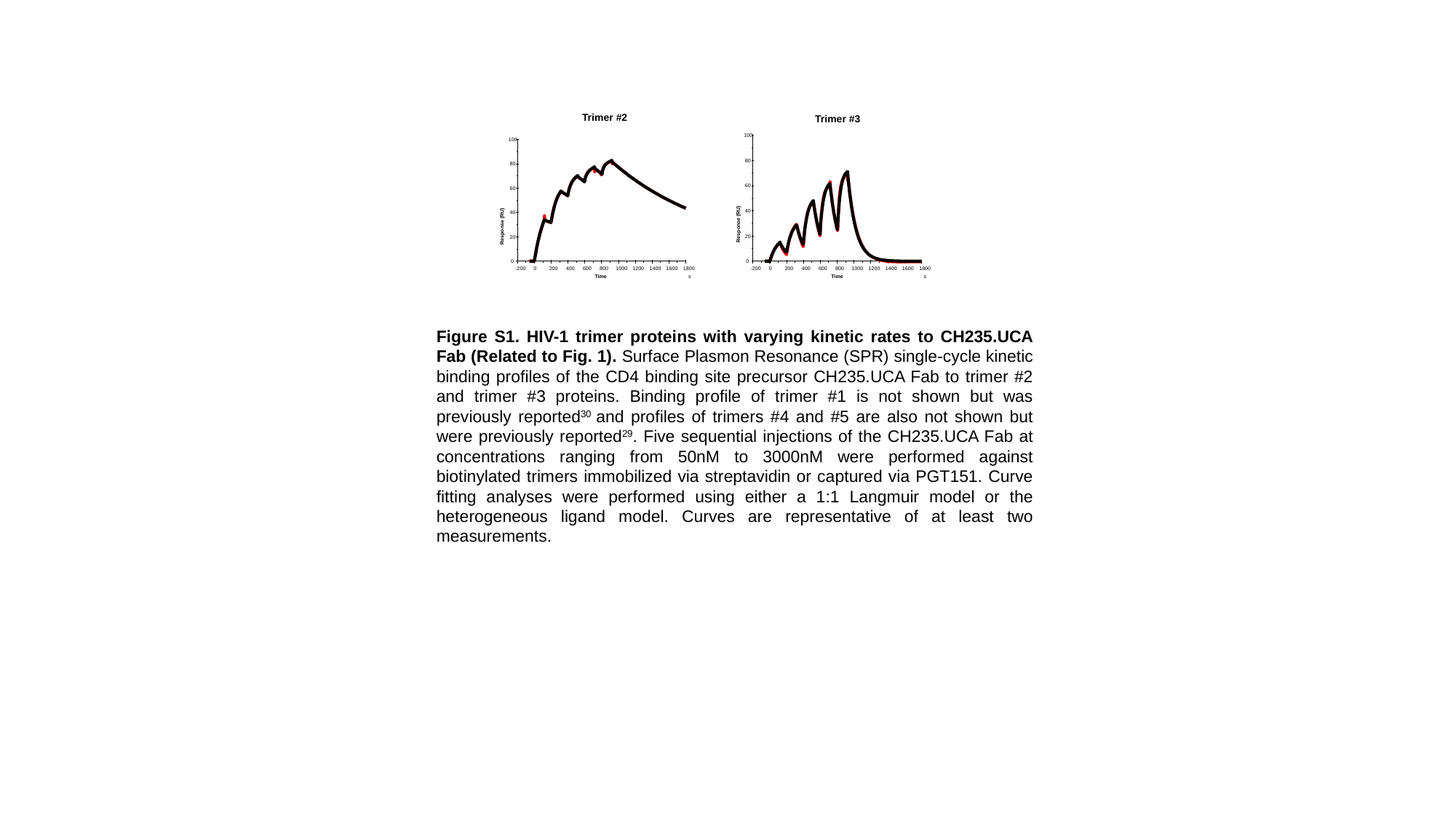

Trimer #2
100
80
60
40
Response (RU)
20
0
-200
0
200
400
600
800
1000
1200
1400
1600
1800
Time
s
Trimer #3
100
80
60
40
Response (RU)
20
0
-200
0
200
400
600
800
1000
1200
1400
1600
1800
Time
s
Figure S1. HIV-1 trimer proteins with varying kinetic rates to CH235.UCA Fab (Related to Fig. 1). Surface Plasmon Resonance (SPR) single-cycle kinetic binding profiles of the CD4 binding site precursor CH235.UCA Fab to trimer #2 and trimer #3 proteins. Binding profile of trimer #1 is not shown but was previously reported^30^ and profiles of trimers #4 and #5 are also not shown but were previously reported^29^. Five sequential injections of the CH235.UCA Fab at concentrations ranging from 50nM to 3000nM were performed against biotinylated trimers immobilized via streptavidin or captured via PGT151. Curve fitting analyses were performed using either a 1:1 Langmuir model or the heterogeneous ligand model. Curves are representative of at least two measurements.

### Slide 3
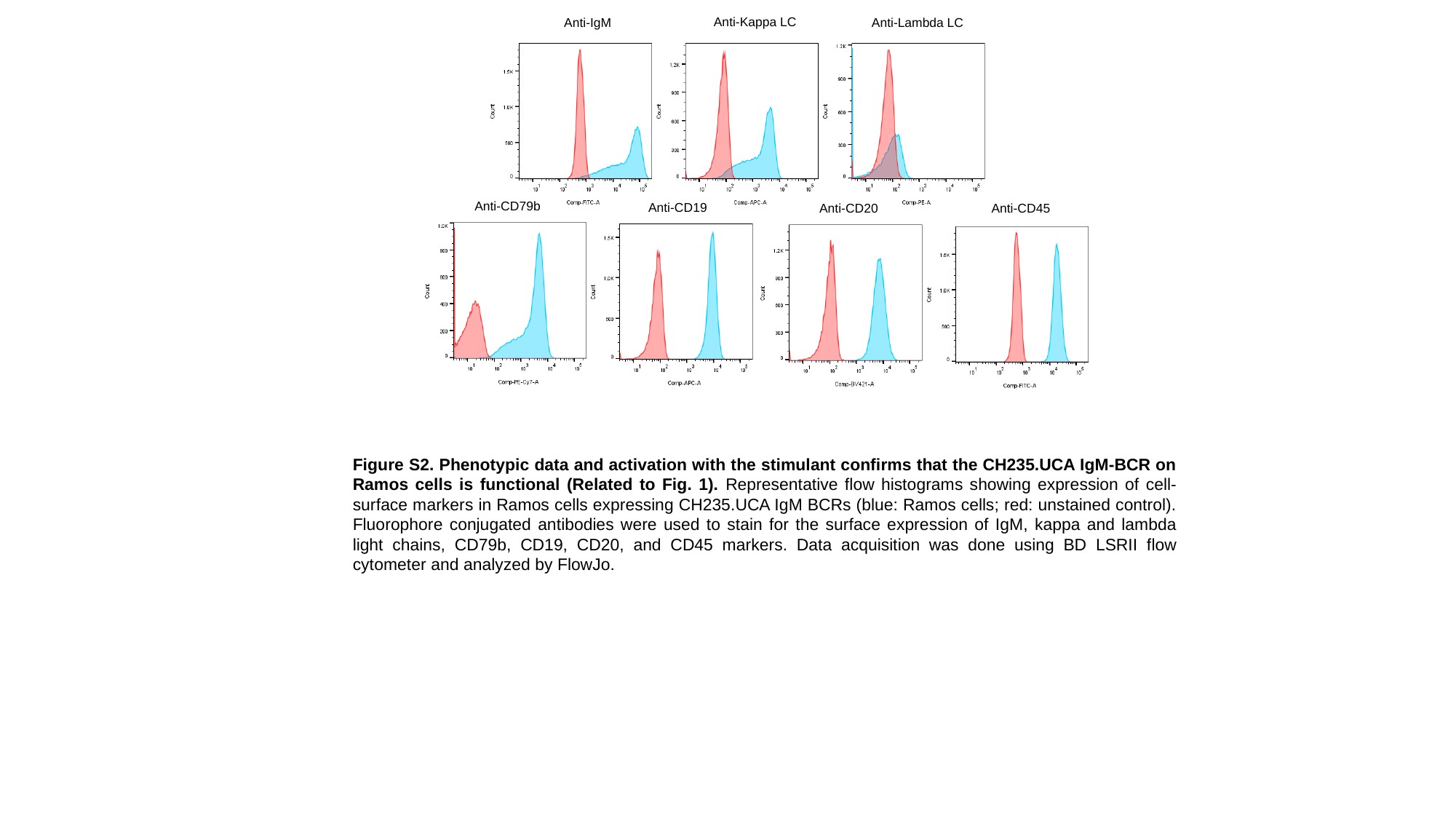

Anti-Kappa LC
Anti-IgM
Anti-Lambda LC
Anti-CD79b
Anti-CD19
Anti-CD20
Anti-CD45
Figure S2. Phenotypic data and activation with the stimulant confirms that the CH235.UCA IgM-BCR on Ramos cells is functional (Related to Fig. 1). Representative flow histograms showing expression of cell-surface markers in Ramos cells expressing CH235.UCA IgM BCRs (blue: Ramos cells; red: unstained control). Fluorophore conjugated antibodies were used to stain for the surface expression of IgM, kappa and lambda light chains, CD79b, CD19, CD20, and CD45 markers. Data acquisition was done using BD LSRII flow cytometer and analyzed by FlowJo.

### Slide 4
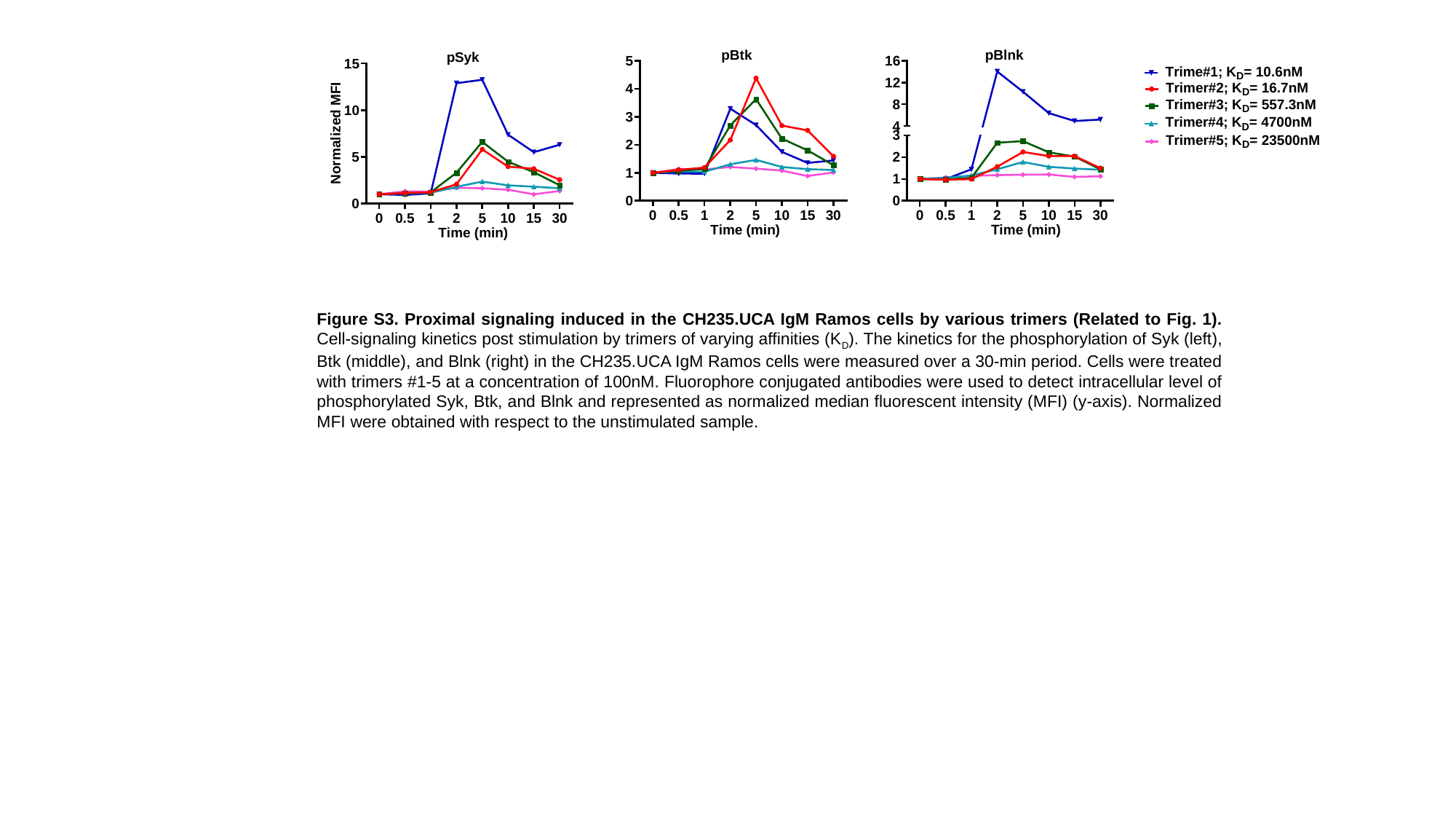

Figure S3. Proximal signaling induced in the CH235.UCA IgM Ramos cells by various trimers (Related to Fig. 1). Cell-signaling kinetics post stimulation by trimers of varying affinities (K_D_). The kinetics for the phosphorylation of Syk (left), Btk (middle), and Blnk (right) in the CH235.UCA IgM Ramos cells were measured over a 30-min period. Cells were treated with trimers #1-5 at a concentration of 100nM. Fluorophore conjugated antibodies were used to detect intracellular level of phosphorylated Syk, Btk, and Blnk and represented as normalized median fluorescent intensity (MFI) (y-axis). Normalized MFI were obtained with respect to the unstimulated sample.

### Slide 5
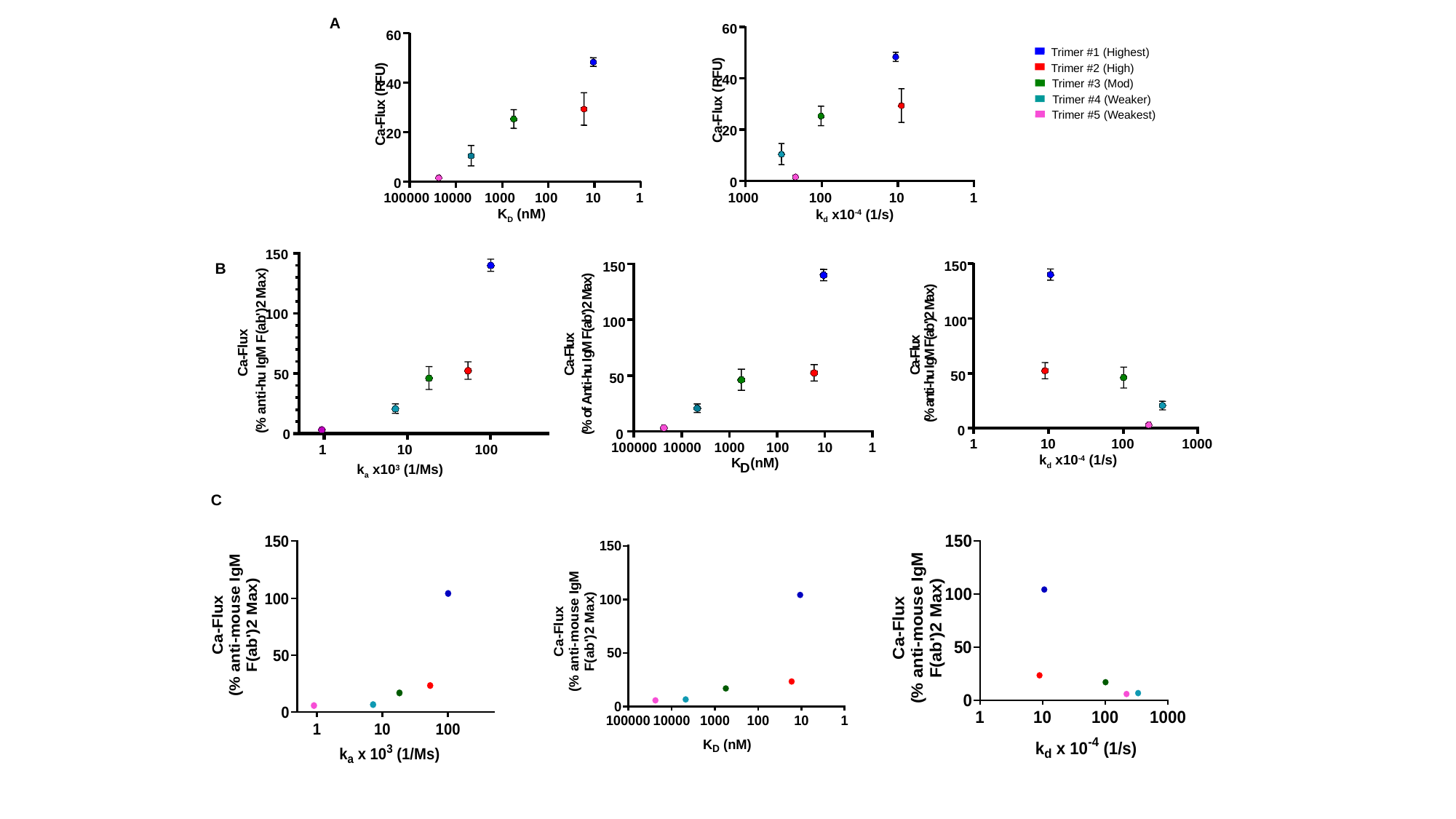

60
)
U
F
40
R
(
x
u
l
F
-
20
a
C
0
1000
100
10
1
k_d_ x10^-4^ (1/s)
A
60
)
U
F
40
R
(
x
u
l
F
-
20
a
C
0
100000
10000
1000
100
10
1
K_D_ (nM)
Trimer #1 (Highest)
Trimer #2 (High)
Trimer #3 (Mod)
Trimer #4 (Weaker)
Trimer #5 (Weakest)
150
100
50
0
1
10
100
k_a_ x10^3^ (1/Ms)
)
x
a
M
2
)
'
b
a
(
x
F
u
l
M
F
-
g
a
I
C
u
h
-
i
t
n
a
%
(
150
100
50
0
100000
10000
1000
100
10
1
K
 (nM)
D
)
x
a
M
2
)
'
b
a
(
F
x
u
M
l
F
g
-
I
a
u
C
h
-
i
t
n
A
f
o
%
(
150
100
50
0
1
10
100
1000
k_d_ x10^-4^ (1/s)
)
x
a
M
2
)
'
b
a
(
x
F
u
l
M
F
-
g
a
I
C
u
h
-
i
t
n
a
%
(
B
C

### Slide 6
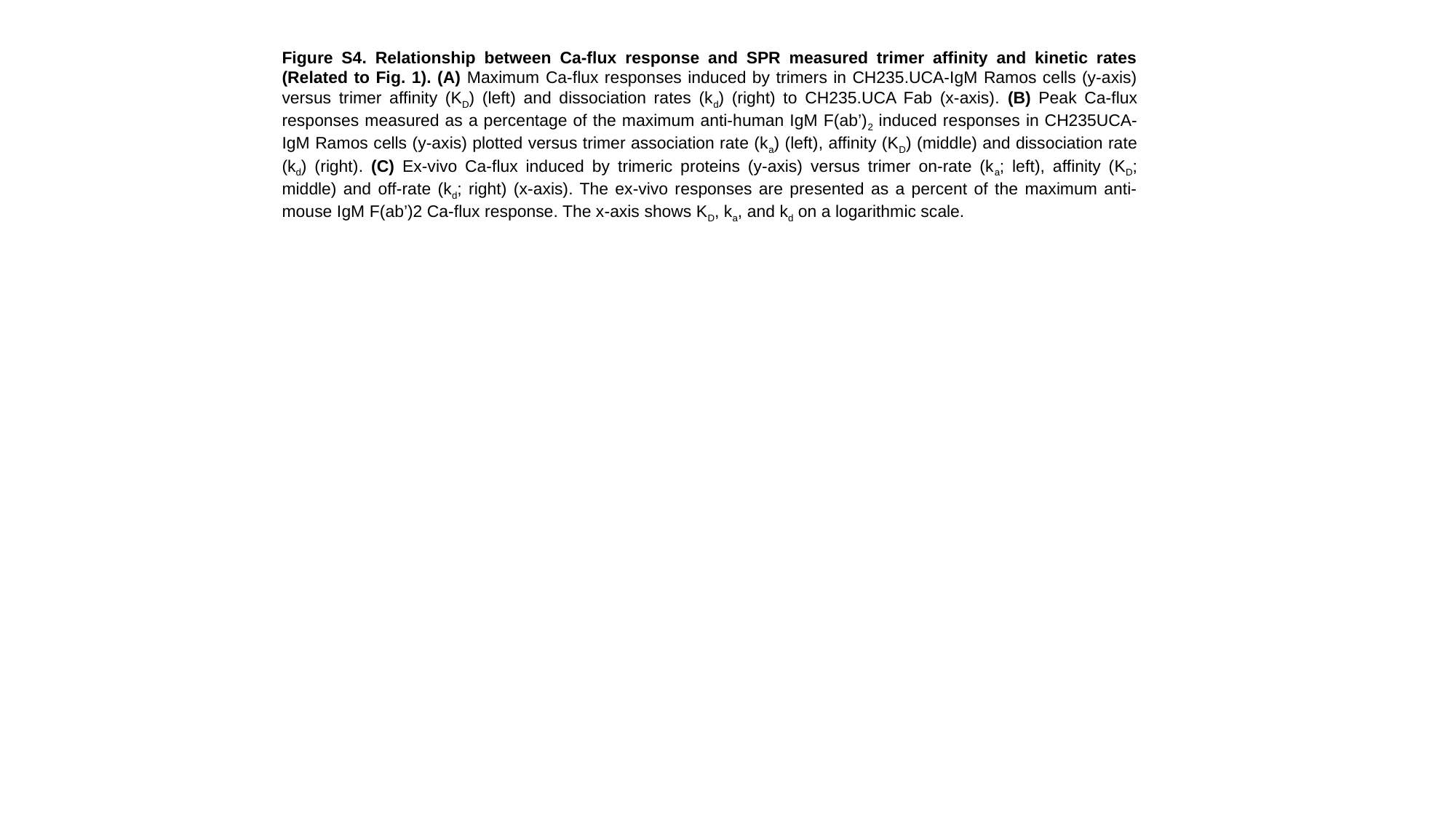

Figure S4. Relationship between Ca-flux response and SPR measured trimer affinity and kinetic rates (Related to Fig. 1). (A) Maximum Ca-flux responses induced by trimers in CH235.UCA-IgM Ramos cells (y-axis) versus trimer affinity (K_D_) (left) and dissociation rates (k_d_) (right) to CH235.UCA Fab (x-axis). (B) Peak Ca-flux responses measured as a percentage of the maximum anti-human IgM F(ab’)_2_ induced responses in CH235UCA-IgM Ramos cells (y-axis) plotted versus trimer association rate (k_a_) (left), affinity (K_D_) (middle) and dissociation rate (k_d_) (right). (C) Ex-vivo Ca-flux induced by trimeric proteins (y-axis) versus trimer on-rate (k_a_; left), affinity (K_D_; middle) and off-rate (k_d_; right) (x-axis). The ex-vivo responses are presented as a percent of the maximum anti-mouse IgM F(ab’)2 Ca-flux response. The x-axis shows K_D_, k_a_, and k_d_ on a logarithmic scale.

### Slide 7
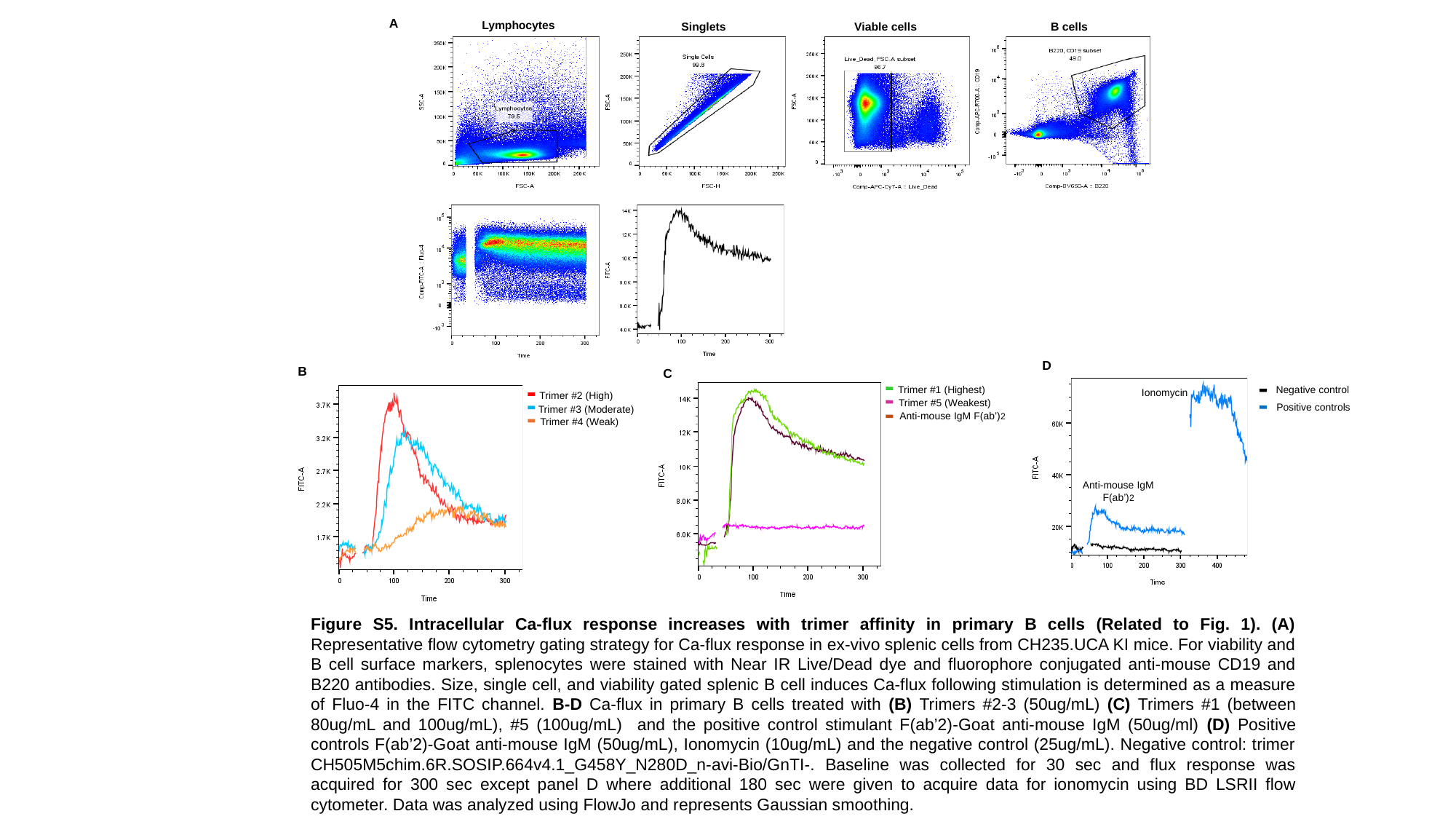

A
Lymphocytes
Singlets
Viable cells
B cells
D
B
C
Trimer #1 (Highest)
Trimer #5 (Weakest)
Anti-mouse IgM F(ab’)2
Negative control
Positive controls
Ionomycin
Trimer #2 (High)
Trimer #3 (Moderate)
Trimer #4 (Weak)
Anti-mouse IgM
F(ab’)2
Figure S5. Intracellular Ca-flux response increases with trimer affinity in primary B cells (Related to Fig. 1). (A) Representative flow cytometry gating strategy for Ca-flux response in ex-vivo splenic cells from CH235.UCA KI mice. For viability and B cell surface markers, splenocytes were stained with Near IR Live/Dead dye and fluorophore conjugated anti-mouse CD19 and B220 antibodies. Size, single cell, and viability gated splenic B cell induces Ca-flux following stimulation is determined as a measure of Fluo-4 in the FITC channel. B-D Ca-flux in primary B cells treated with (B) Trimers #2-3 (50ug/mL) (C) Trimers #1 (between 80ug/mL and 100ug/mL), #5 (100ug/mL) and the positive control stimulant F(ab’2)-Goat anti-mouse IgM (50ug/ml) (D) Positive controls F(ab’2)-Goat anti-mouse IgM (50ug/mL), Ionomycin (10ug/mL) and the negative control (25ug/mL). Negative control: trimer CH505M5chim.6R.SOSIP.664v4.1_G458Y_N280D_n-avi-Bio/GnTI-. Baseline was collected for 30 sec and flux response was acquired for 300 sec except panel D where additional 180 sec were given to acquire data for ionomycin using BD LSRII flow cytometer. Data was analyzed using FlowJo and represents Gaussian smoothing.

### Slide 8
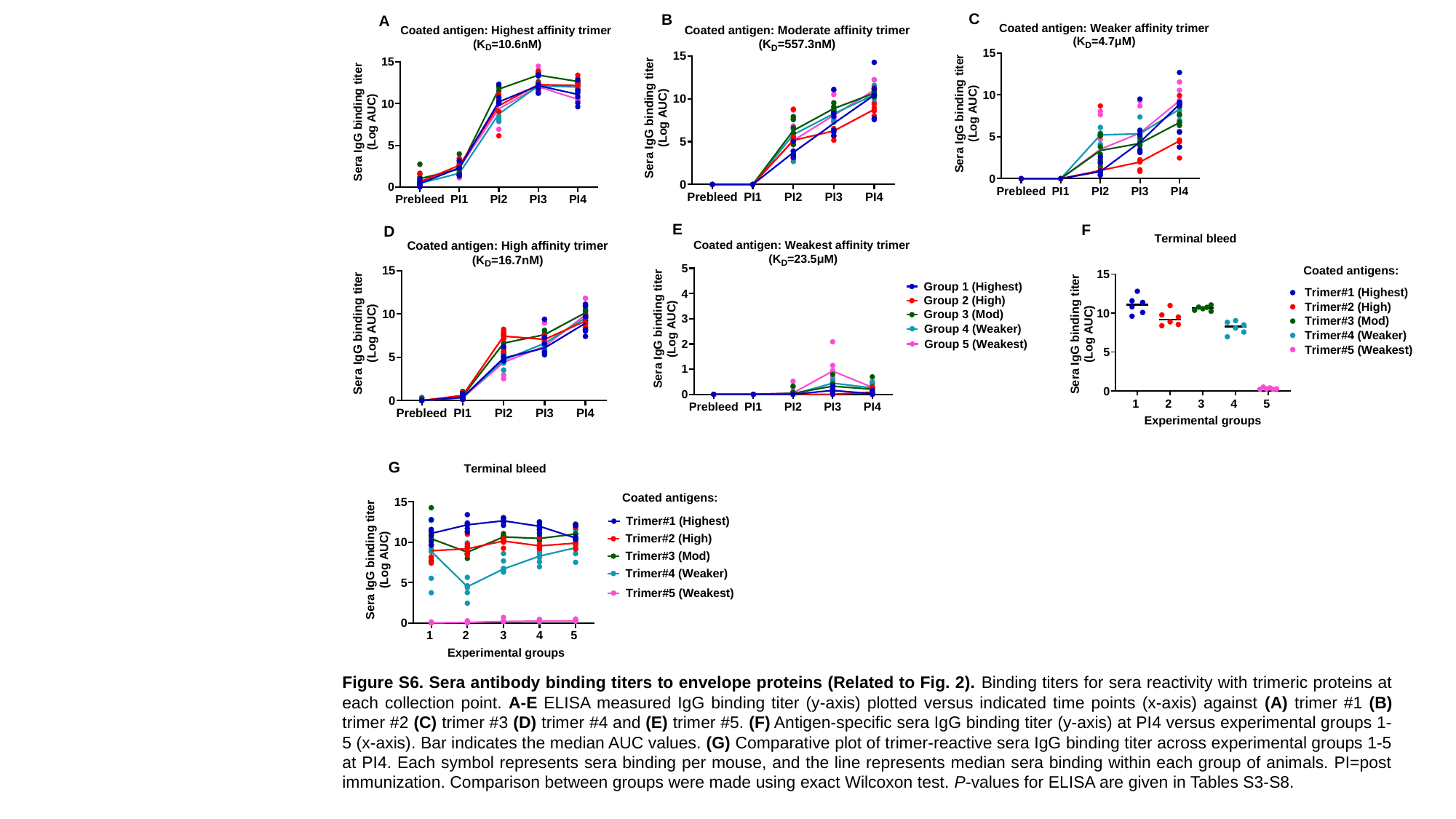

C
B
A
E
F
D
G
Figure S6. Sera antibody binding titers to envelope proteins (Related to Fig. 2). Binding titers for sera reactivity with trimeric proteins at each collection point. A-E ELISA measured IgG binding titer (y-axis) plotted versus indicated time points (x-axis) against (A) trimer #1 (B) trimer #2 (C) trimer #3 (D) trimer #4 and (E) trimer #5. (F) Antigen-specific sera IgG binding titer (y-axis) at PI4 versus experimental groups 1-5 (x-axis). Bar indicates the median AUC values. (G) Comparative plot of trimer-reactive sera IgG binding titer across experimental groups 1-5 at PI4. Each symbol represents sera binding per mouse, and the line represents median sera binding within each group of animals. PI=post immunization. Comparison between groups were made using exact Wilcoxon test. P-values for ELISA are given in Tables S3-S8.

### Slide 9
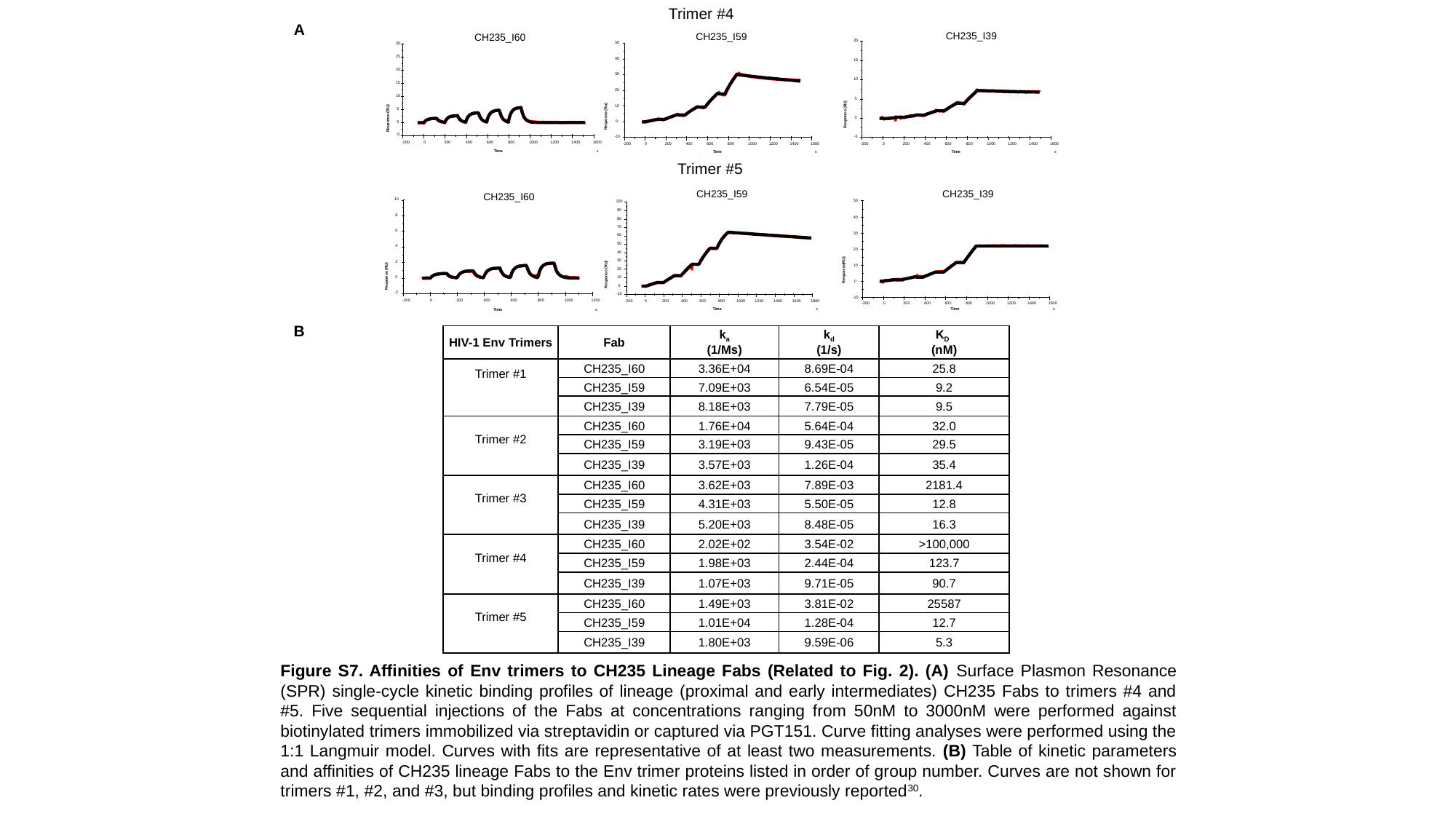

Trimer #4
A
CH235_I39
20
15
10
5
Response (RU)
0
-5
-200
0
200
400
600
800
1000
1200
1400
1600
Time
s
CH235_I59
50
40
30
20
10
Response (Ru)
0
-10
-200
0
200
400
600
800
1000
1200
1400
1600
Time
s
CH235_I60
30
25
20
15
10
5
Response (RU)
0
-5
-200
0
200
400
600
800
1000
1200
1400
1600
Time
s
Trimer #5
Trimer #5
CH235_I59
100
90
80
70
60
50
40
30
20
Response (RU)
10
0
-10
-200
0
200
400
600
800
1000
1200
1400
1600
1800
Time
s
CH235_I39
50
40
30
20
10
Response(RU)
0
-10
-200
0
200
400
600
800
1000
1200
1400
1600
Time
s
CH235_I60
10
8
6
4
2
Response (RU)
0
-2
-200
0
200
400
600
800
1000
1200
Time
s
B
| HIV-1 Env Trimers | Fab | k_a_ (1/Ms) | k_d_ (1/s) | K_D_ (nM) |
| --- | --- | --- | --- | --- |
| Trimer #1 | CH235\_I60 | 3.36E+04 | 8.69E-04 | 25.8 |
| | CH235\_I59 | 7.09E+03 | 6.54E-05 | 9.2 |
| | CH235\_I39 | 8.18E+03 | 7.79E-05 | 9.5 |
| Trimer #2 | CH235\_I60 | 1.76E+04 | 5.64E-04 | 32.0 |
| | CH235\_I59 | 3.19E+03 | 9.43E-05 | 29.5 |
| | CH235\_I39 | 3.57E+03 | 1.26E-04 | 35.4 |
| Trimer #3 | CH235\_I60 | 3.62E+03 | 7.89E-03 | 2181.4 |
| | CH235\_I59 | 4.31E+03 | 5.50E-05 | 12.8 |
| | CH235\_I39 | 5.20E+03 | 8.48E-05 | 16.3 |
| Trimer #4 | CH235\_I60 | 2.02E+02 | 3.54E-02 | >100,000 |
| | CH235\_I59 | 1.98E+03 | 2.44E-04 | 123.7 |
| | CH235\_I39 | 1.07E+03 | 9.71E-05 | 90.7 |
| Trimer #5 | CH235\_I60 | 1.49E+03 | 3.81E-02 | 25587 |
| | CH235\_I59 | 1.01E+04 | 1.28E-04 | 12.7 |
| | CH235\_I39 | 1.80E+03 | 9.59E-06 | 5.3 |
Figure S7. Affinities of Env trimers to CH235 Lineage Fabs (Related to Fig. 2). (A) Surface Plasmon Resonance (SPR) single-cycle kinetic binding profiles of lineage (proximal and early intermediates) CH235 Fabs to trimers #4 and #5. Five sequential injections of the Fabs at concentrations ranging from 50nM to 3000nM were performed against biotinylated trimers immobilized via streptavidin or captured via PGT151. Curve fitting analyses were performed using the 1:1 Langmuir model. Curves with fits are representative of at least two measurements. (B) Table of kinetic parameters and affinities of CH235 lineage Fabs to the Env trimer proteins listed in order of group number. Curves are not shown for trimers #1, #2, and #3, but binding profiles and kinetic rates were previously reported^30^.

### Slide 10
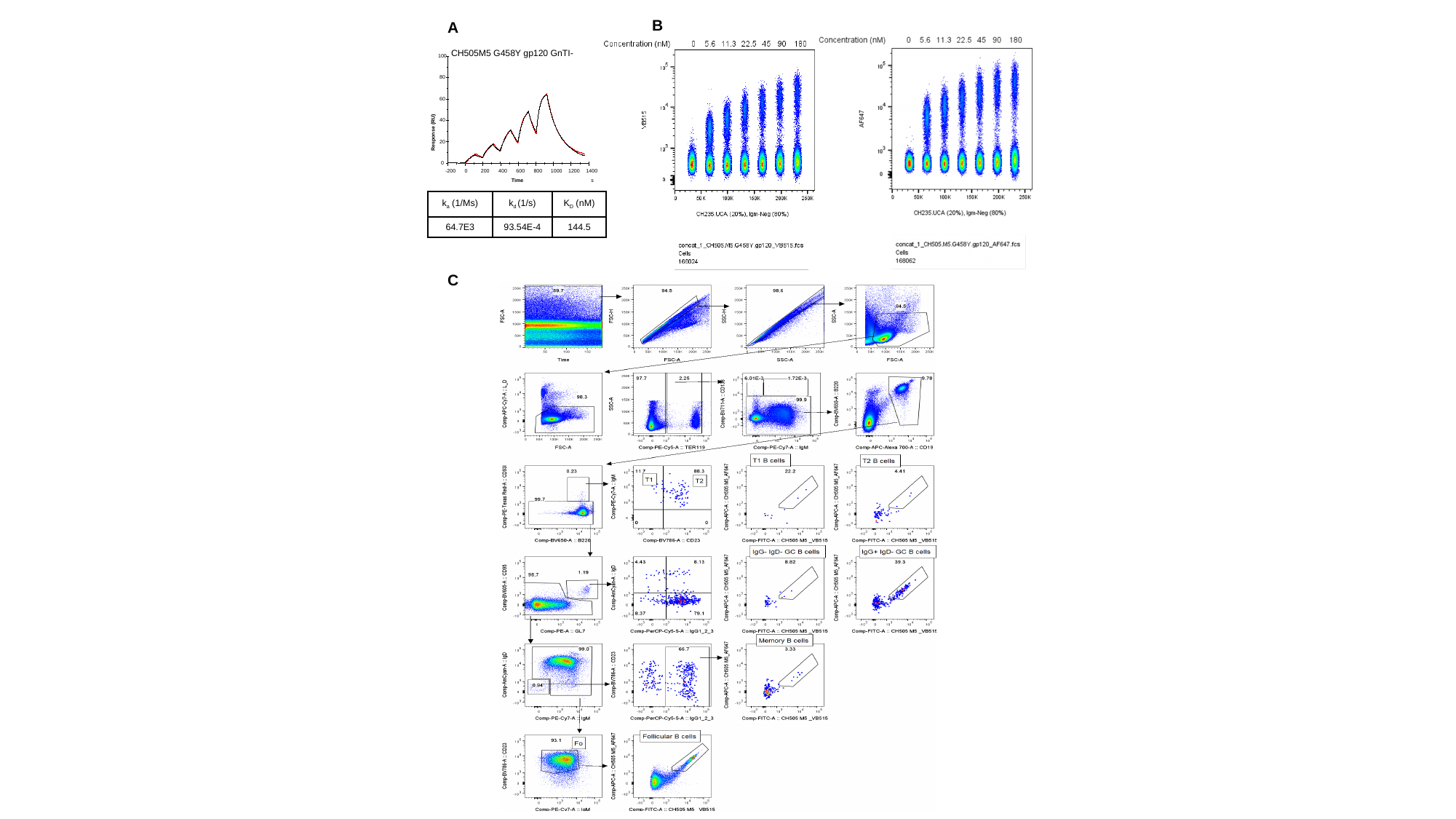

B
A
CH505M5 G458Y gp120 GnTI-
100
80
60
40
Response (RU)
20
0
-200
0
200
400
600
800
1000
1200
1400
Time
s
| k_a_ (1/Ms) | k_d_ (1/s) | K_D_ (nM) |
| --- | --- | --- |
| 64.7E3 | 93.54E-4 | 144.5 |
C

### Slide 11
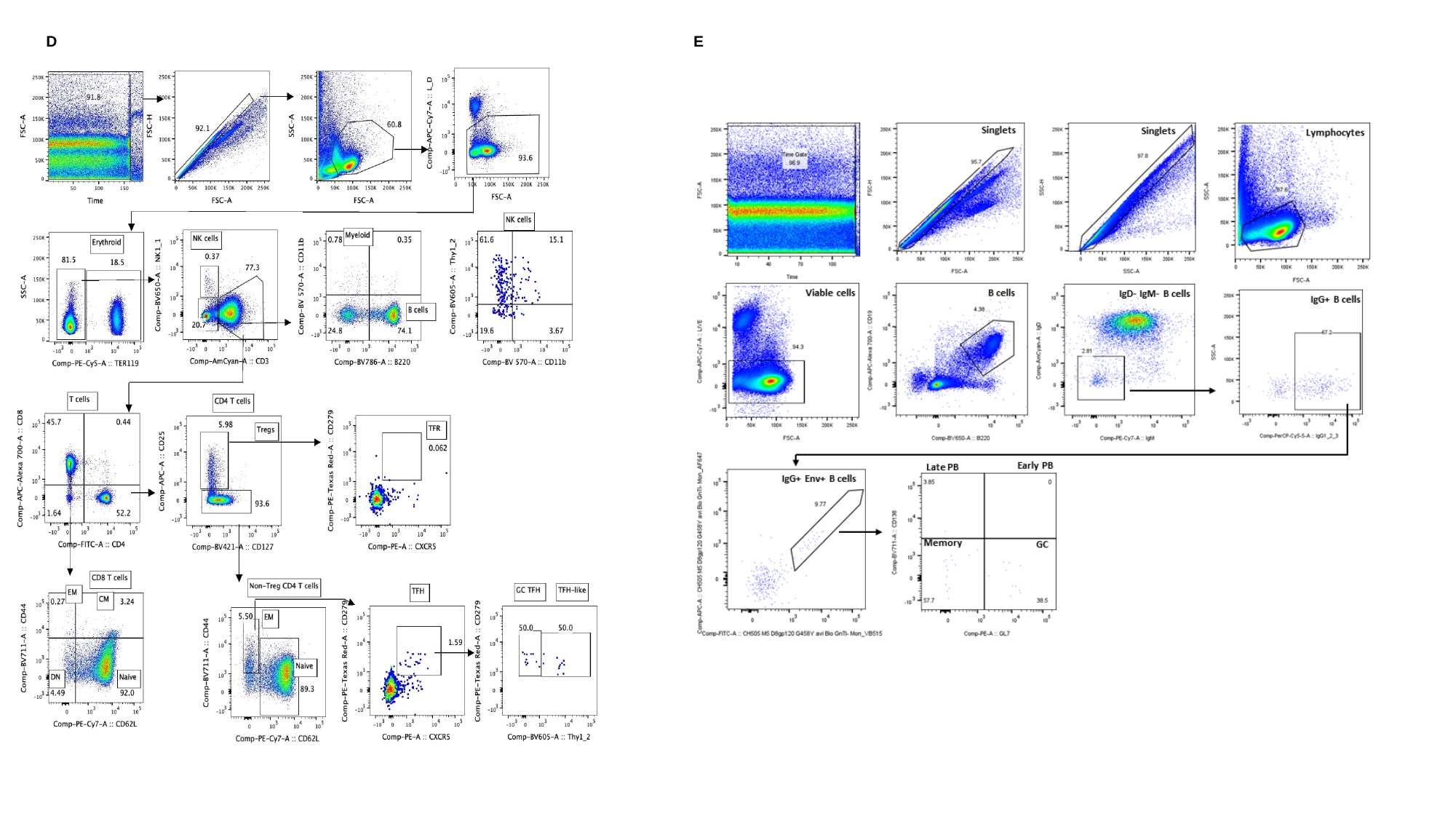

D
E

### Slide 12
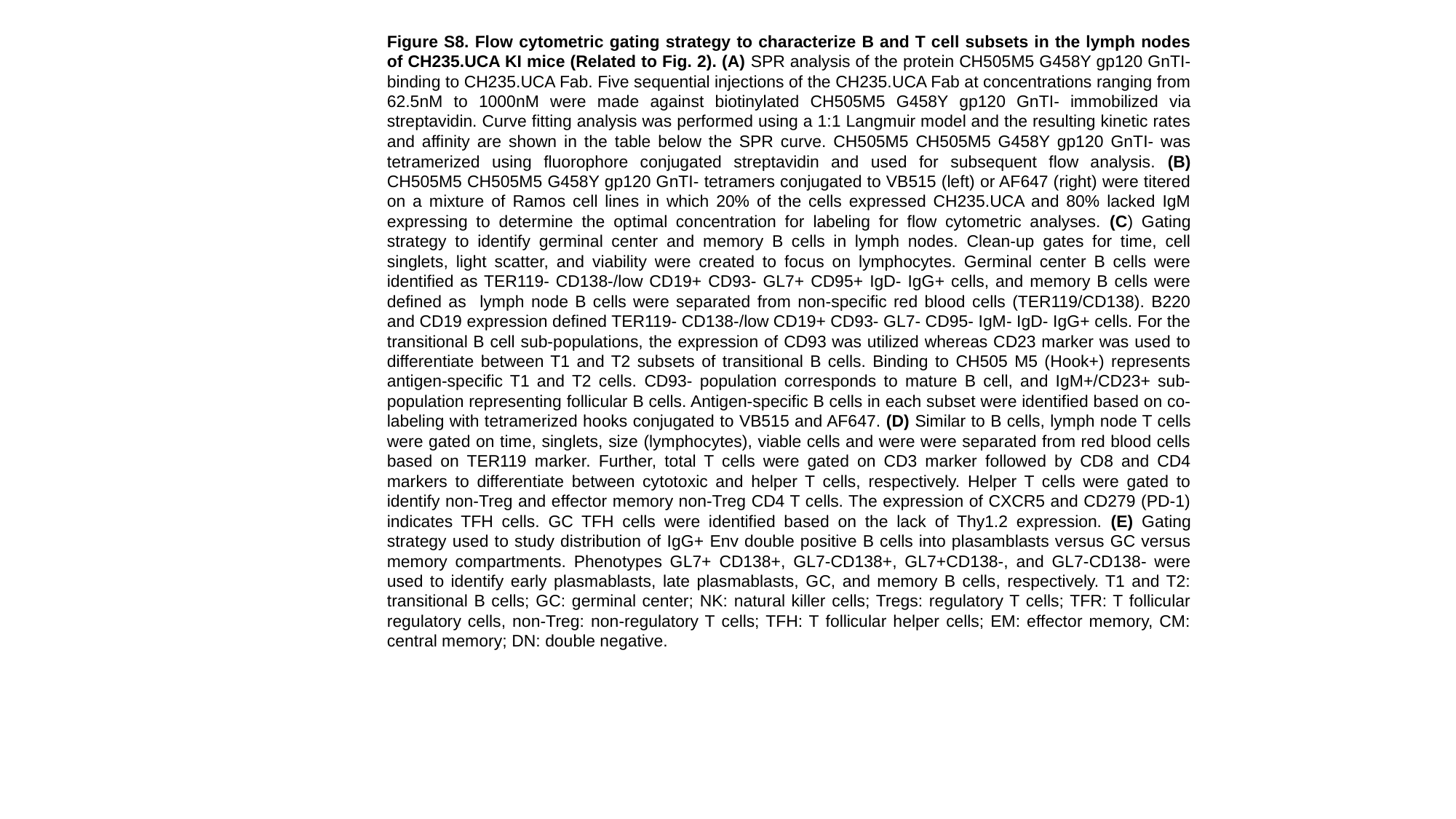

Figure S8. Flow cytometric gating strategy to characterize B and T cell subsets in the lymph nodes of CH235.UCA KI mice (Related to Fig. 2). (A) SPR analysis of the protein CH505M5 G458Y gp120 GnTI- binding to CH235.UCA Fab. Five sequential injections of the CH235.UCA Fab at concentrations ranging from 62.5nM to 1000nM were made against biotinylated CH505M5 G458Y gp120 GnTI- immobilized via streptavidin. Curve fitting analysis was performed using a 1:1 Langmuir model and the resulting kinetic rates and affinity are shown in the table below the SPR curve. CH505M5 CH505M5 G458Y gp120 GnTI- was tetramerized using fluorophore conjugated streptavidin and used for subsequent flow analysis. (B) CH505M5 CH505M5 G458Y gp120 GnTI- tetramers conjugated to VB515 (left) or AF647 (right) were titered on a mixture of Ramos cell lines in which 20% of the cells expressed CH235.UCA and 80% lacked IgM expressing to determine the optimal concentration for labeling for flow cytometric analyses. (C) Gating strategy to identify germinal center and memory B cells in lymph nodes. Clean-up gates for time, cell singlets, light scatter, and viability were created to focus on lymphocytes. Germinal center B cells were identified as TER119- CD138-/low CD19+ CD93- GL7+ CD95+ IgD- IgG+ cells, and memory B cells were defined as lymph node B cells were separated from non-specific red blood cells (TER119/CD138). B220 and CD19 expression defined TER119- CD138-/low CD19+ CD93- GL7- CD95- IgM- IgD- IgG+ cells. For the transitional B cell sub-populations, the expression of CD93 was utilized whereas CD23 marker was used to differentiate between T1 and T2 subsets of transitional B cells. Binding to CH505 M5 (Hook+) represents antigen-specific T1 and T2 cells. CD93- population corresponds to mature B cell, and IgM+/CD23+ sub-population representing follicular B cells. Antigen-specific B cells in each subset were identified based on co-labeling with tetramerized hooks conjugated to VB515 and AF647. (D) Similar to B cells, lymph node T cells were gated on time, singlets, size (lymphocytes), viable cells and were were separated from red blood cells based on TER119 marker. Further, total T cells were gated on CD3 marker followed by CD8 and CD4 markers to differentiate between cytotoxic and helper T cells, respectively. Helper T cells were gated to identify non-Treg and effector memory non-Treg CD4 T cells. The expression of CXCR5 and CD279 (PD-1) indicates TFH cells. GC TFH cells were identified based on the lack of Thy1.2 expression. (E) Gating strategy used to study distribution of IgG+ Env double positive B cells into plasamblasts versus GC versus memory compartments. Phenotypes GL7+ CD138+, GL7-CD138+, GL7+CD138-, and GL7-CD138- were used to identify early plasmablasts, late plasmablasts, GC, and memory B cells, respectively. T1 and T2: transitional B cells; GC: germinal center; NK: natural killer cells; Tregs: regulatory T cells; TFR: T follicular regulatory cells, non-Treg: non-regulatory T cells; TFH: T follicular helper cells; EM: effector memory, CM: central memory; DN: double negative.

### Slide 13
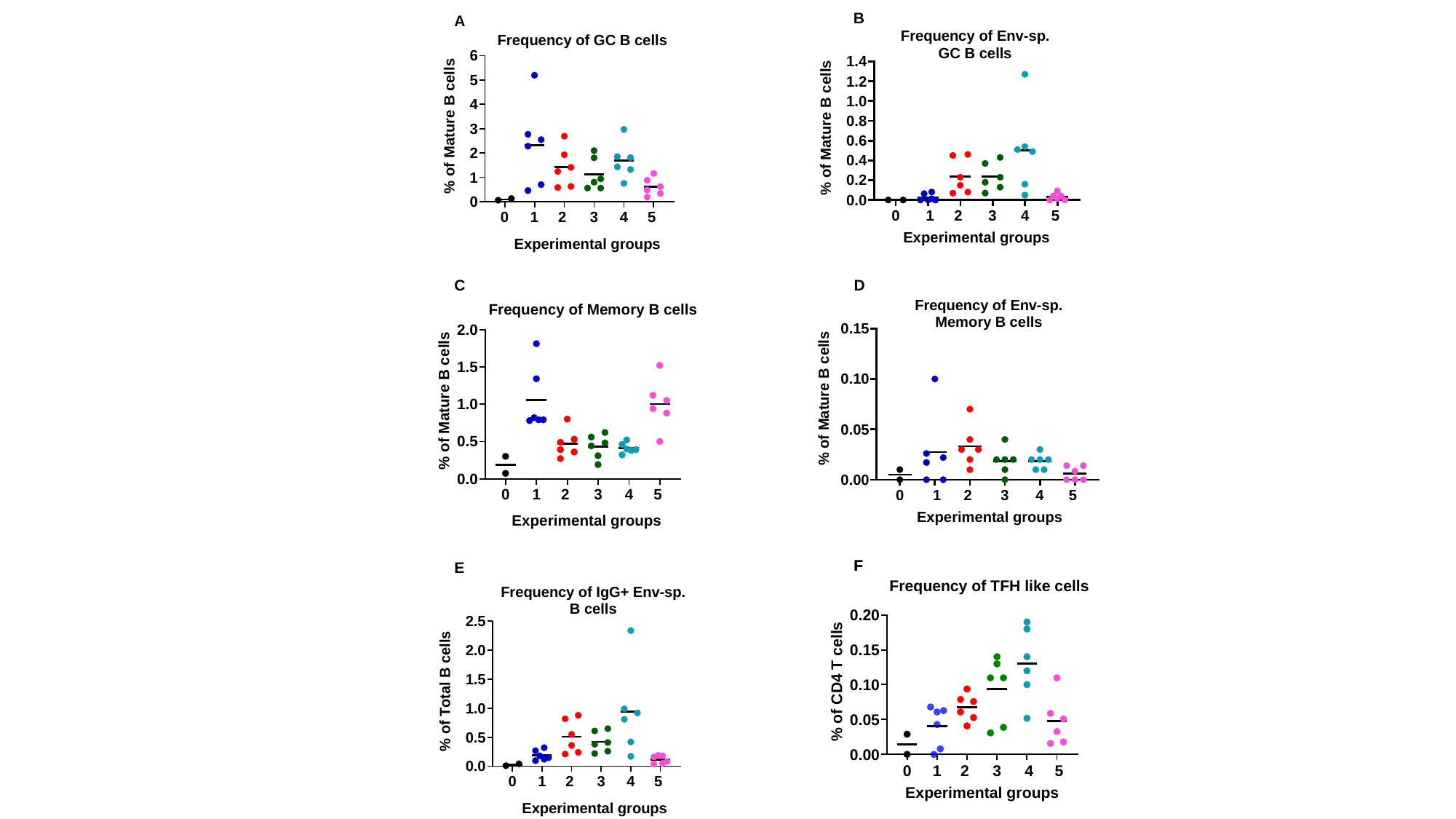

B
A
C
D
F
F
E

### Slide 14
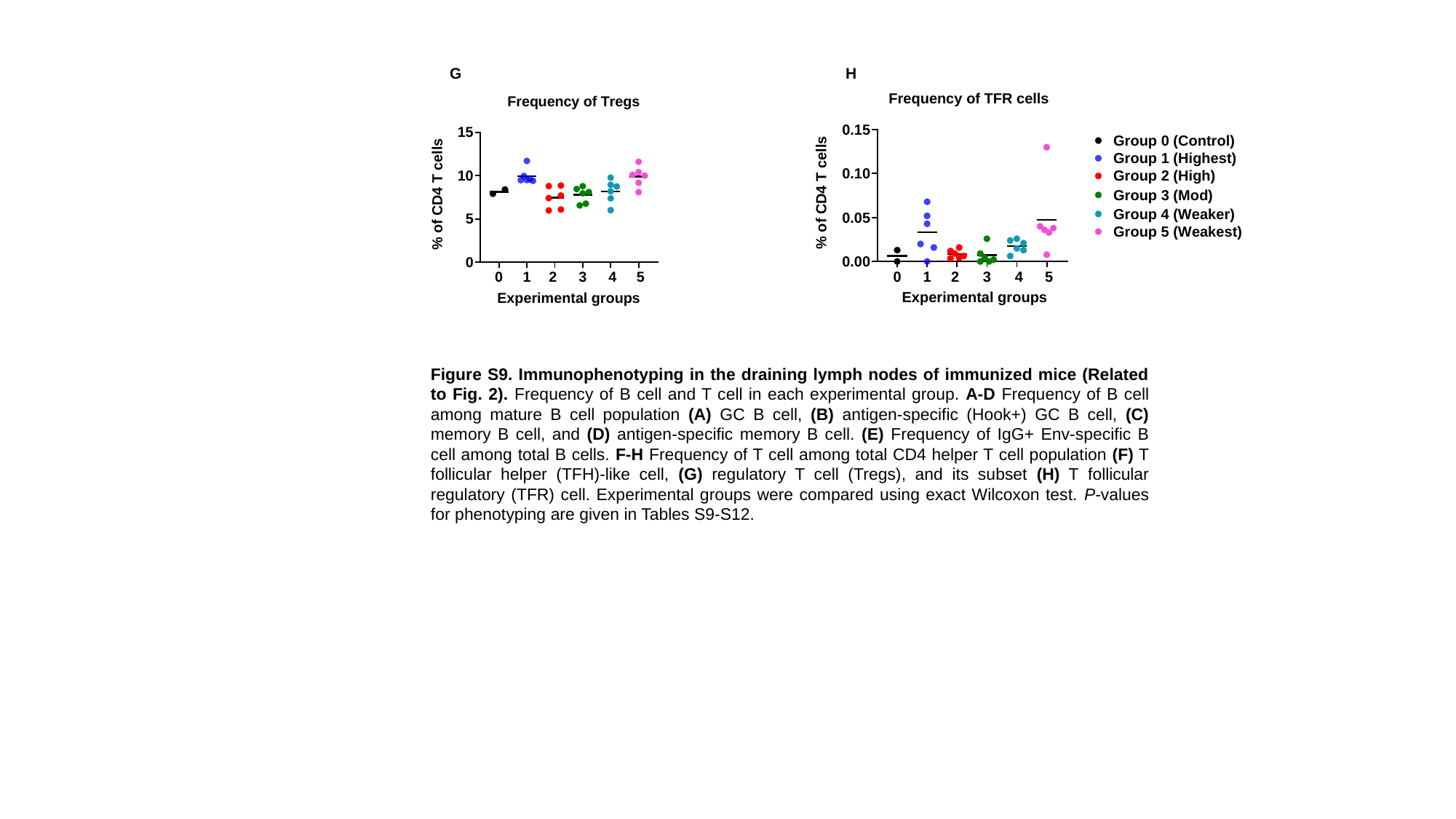

G
H
Figure S9. Immunophenotyping in the draining lymph nodes of immunized mice (Related to Fig. 2). Frequency of B cell and T cell in each experimental group. A-D Frequency of B cell among mature B cell population (A) GC B cell, (B) antigen-specific (Hook+) GC B cell, (C) memory B cell, and (D) antigen-specific memory B cell. (E) Frequency of IgG+ Env-specific B cell among total B cells. F-H Frequency of T cell among total CD4 helper T cell population (F) T follicular helper (TFH)-like cell, (G) regulatory T cell (Tregs), and its subset (H) T follicular regulatory (TFR) cell. Experimental groups were compared using exact Wilcoxon test. P-values for phenotyping are given in Tables S9-S12.

### Slide 15
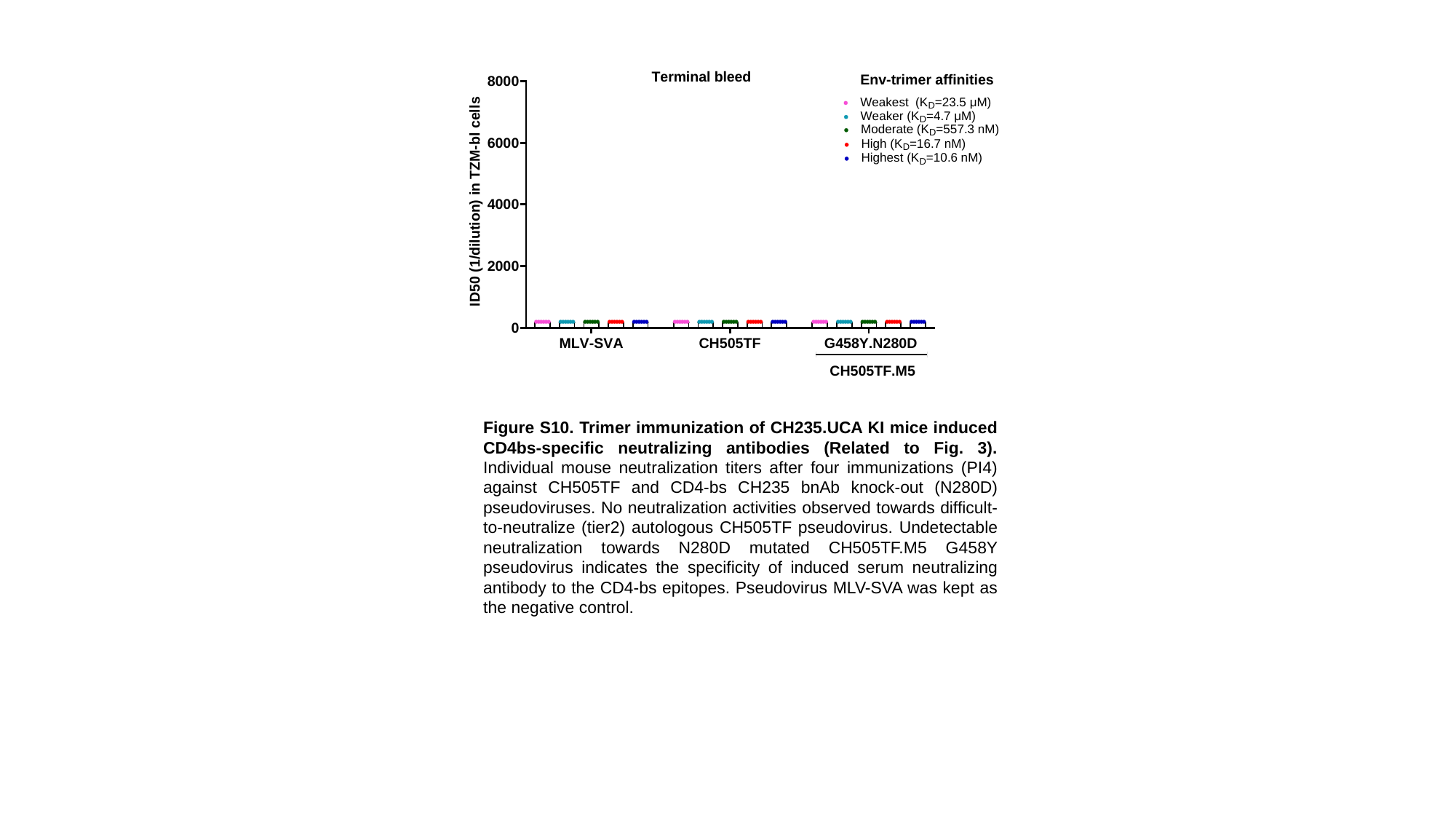

Figure S10. Trimer immunization of CH235.UCA KI mice induced CD4bs-specific neutralizing antibodies (Related to Fig. 3). Individual mouse neutralization titers after four immunizations (PI4) against CH505TF and CD4-bs CH235 bnAb knock-out (N280D) pseudoviruses. No neutralization activities observed towards difficult-to-neutralize (tier2) autologous CH505TF pseudovirus. Undetectable neutralization towards N280D mutated CH505TF.M5 G458Y pseudovirus indicates the specificity of induced serum neutralizing antibody to the CD4-bs epitopes. Pseudovirus MLV-SVA was kept as the negative control.

### Slide 16
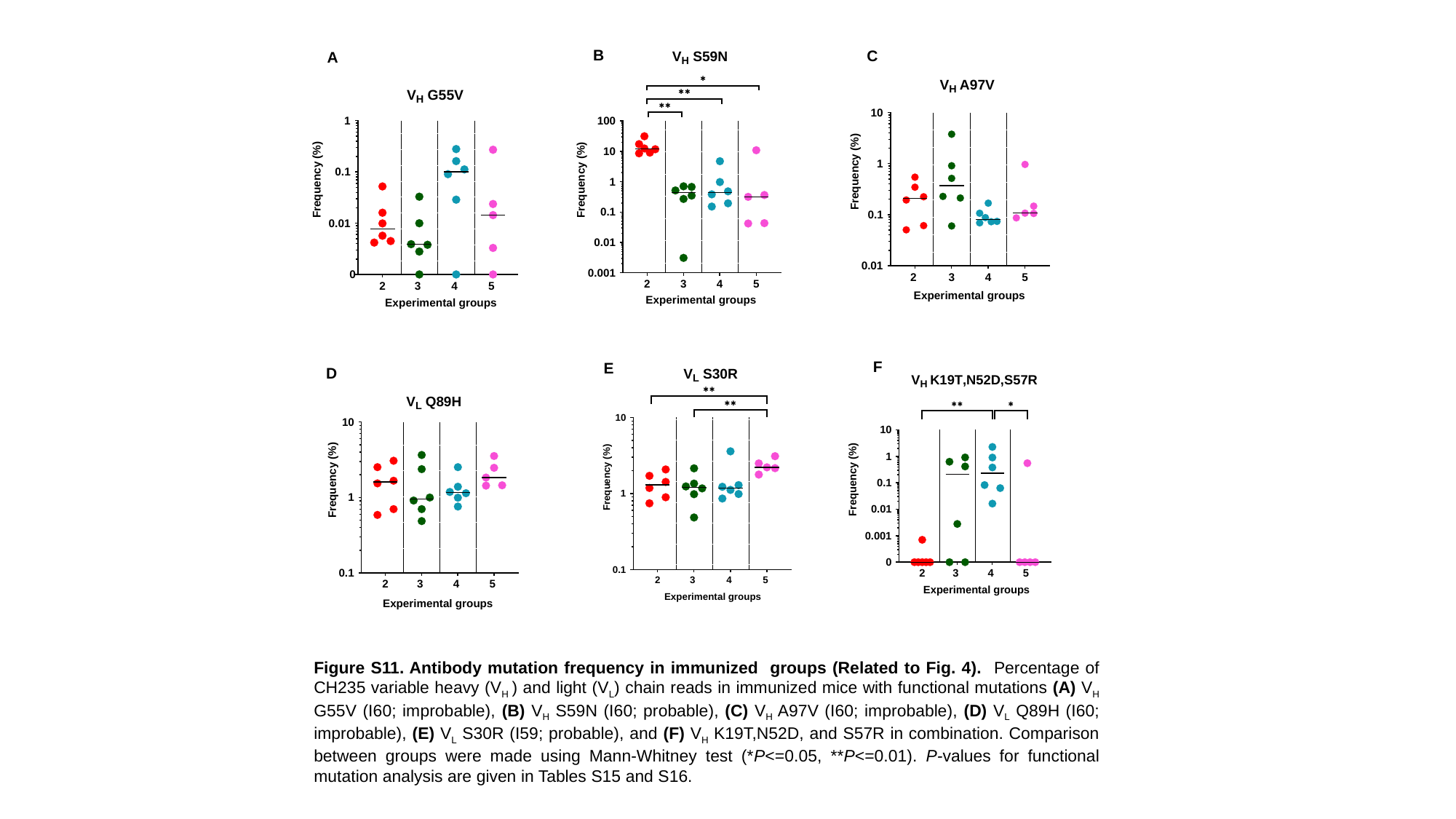

B
C
A
F
E
D
Figure S11. Antibody mutation frequency in immunized groups (Related to Fig. 4). Percentage of CH235 variable heavy (V_H_ ) and light (V_L_) chain reads in immunized mice with functional mutations (A) V_H_ G55V (I60; improbable), (B) V_H_ S59N (I60; probable), (C) V_H_ A97V (I60; improbable), (D) V_L_ Q89H (I60; improbable), (E) V_L_ S30R (I59; probable), and (F) V_H_ K19T,N52D, and S57R in combination. Comparison between groups were made using Mann-Whitney test (*P<=0.05, **P<=0.01). P-values for functional mutation analysis are given in Tables S15 and S16.

### Slide 17
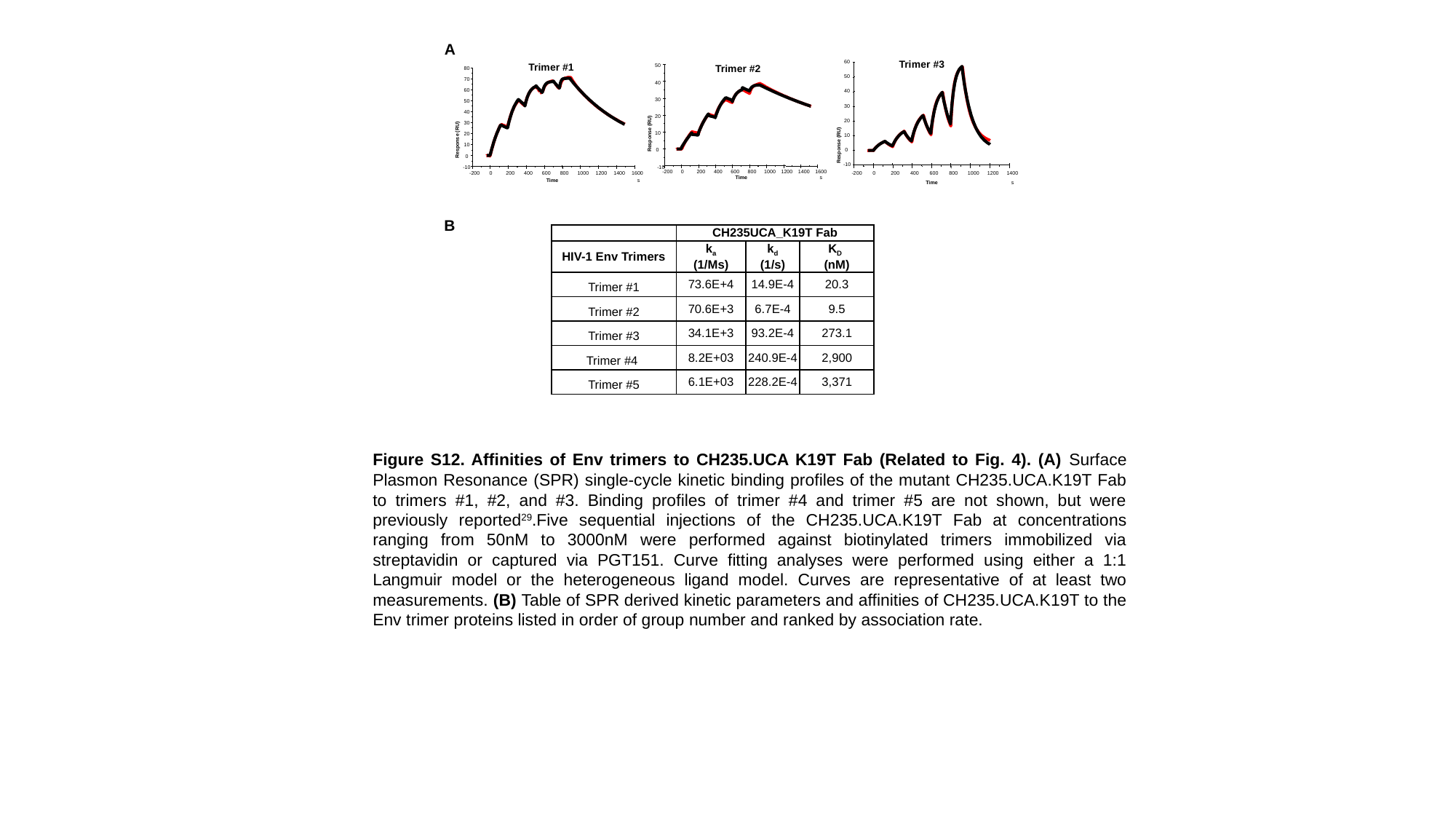

A
60
50
40
30
20
10
Response (RU)
0
-10
-200
0
200
400
600
800
1000
1200
1400
Time
s
Trimer #3
Trimer #1
50
40
30
20
10
Response (RU)
0
-10
-200
0
200
400
600
800
1000
1200
1400
1600
Time
s
Trimer #2
80
70
60
50
40
30
20
Response (RU)
10
0
-10
-200
0
200
400
600
800
1000
1200
1400
1600
Time
s
B
| | CH235UCA\_K19T Fab | | |
| --- | --- | --- | --- |
| HIV-1 Env Trimers | k_a_ (1/Ms) | k_d_ (1/s) | K_D_ (nM) |
| Trimer #1 | 73.6E+4 | 14.9E-4 | 20.3 |
| Trimer #2 | 70.6E+3 | 6.7E-4 | 9.5 |
| Trimer #3 | 34.1E+3 | 93.2E-4 | 273.1 |
| Trimer #4 | 8.2E+03 | 240.9E-4 | 2,900 |
| Trimer #5 | 6.1E+03 | 228.2E-4 | 3,371 |
Figure S12. Affinities of Env trimers to CH235.UCA K19T Fab (Related to Fig. 4). (A) Surface Plasmon Resonance (SPR) single-cycle kinetic binding profiles of the mutant CH235.UCA.K19T Fab to trimers #1, #2, and #3. Binding profiles of trimer #4 and trimer #5 are not shown, but were previously reported^29^.Five sequential injections of the CH235.UCA.K19T Fab at concentrations ranging from 50nM to 3000nM were performed against biotinylated trimers immobilized via streptavidin or captured via PGT151. Curve fitting analyses were performed using either a 1:1 Langmuir model or the heterogeneous ligand model. Curves are representative of at least two measurements. (B) Table of SPR derived kinetic parameters and affinities of CH235.UCA.K19T to the Env trimer proteins listed in order of group number and ranked by association rate.
